# Abnormal hyperactivity of specific striatal ensembles encodes distinct dyskinetic behaviors revealed by high-resolution clustering

**DOI:** 10.1101/2024.09.06.611664

**Authors:** Cristina Alcacer, Andreas Klaus, Marcelo Mendonça, Sara F. Abalde, Maria Angela Cenci, Rui M. Costa

## Abstract

L-DOPA-induced dyskinesia (LID) is a debilitating complication of dopamine replacement therapy in Parkinsońs disease and the most common hyperkinetic disorder of basal ganglia origin. Abnormal activity of striatal D1 and D2 spiny projection neurons (SPNs) is critical for LID, yet the link between SPN activity patterns and specific dyskinetic movements remains unknown. To explore this, we developed a novel method for clustering movements based on high-resolution motion sensors and video recordings. In a mouse model of LID, this method identified two main dyskinesia types and pathological rotations, all absent during normal behavior. Using single-cell resolution imaging, we found that specific sets of both D1 and D2-SPNs were abnormally active during these pathological movements. Under baseline conditions, the same SPN sets were active during behaviors sharing physical features with LID movements. These findings indicate that ensembles of behavior-encoding D1- and D2-SPNs form new combinations of hyperactive neurons mediating specific dyskinetic movements.

## Introduction

In Parkinson’s disease (PD), the degeneration of dopaminergic neurons that project to the striatum causes poverty and slowness of movement. Dopamine (DA) replacement therapy with L-DOPA is still the most effective treatment, but leads to a development of abnormal involuntary movements in the majority of patients within a few years ^1^. Although the involuntary movements are collectively referred to as L-DOPA-induced dyskinesia (LID), they have variable clinical presentations, manifesting with different combinations of fast hyperkinetic motions and dystonic features (sustained twisting movements and abnormal postures) in different body parts ^2^. In addition to being a medically important problem, LID provides a study paradigm to unveil patterns of striatal activity disrupting the control of movement sequences ^3–6^.

Movement control depends on the interplay of two main pathways originating from two populations of striatal neurons. Spiny projection neurons (SPN) expressing DA D1-receptor (D1-SPNs) project directly to the basal ganglia output nuclei forming the classically called ‘direct pathway’, while D2-receptor expressing SPNs (D2-SPNs) influence basal ganglia output indirectly via intermediate nuclei ^7,8^. Canonical models postulate that direct and indirect pathways have opposite effects on movement: activation of D1-SPN facilitates movement whereas activation of D2-SPN inhibits movement ^9–11^. However, accumulating evidence suggests that both pathways increase their activity at movement onset and that coordinated activation of D1- and D2-SPNs is required for proper action initiation ^12–15^. The role of SPNs in LID has been studied using parkinsonian rodent models treated with L-DOPA and developing axial, limb, and orofacial abnormal involuntary movements (AIMs). The recording of striatal activity during the expression of AIMs has revealed opposite changes in the average activity levels of D1-SPNs and D2-SPNs ^5,6^. Moreover, SPN type-specific stimulations using chemogenetic ^9^ or optogenetic methods ^4,6,16^ concordantly show that LID is aggravated by increasing the activity of D1-SPNs and blunted by stimulating D2-SPNs. Taken together, these studies have established a causal link between LID and a disrupted interplay between direct and indirect pathway, with concomitant D1-SPN hyperactivity and D2-SPN underactivity. However, this level of explanation cannot account for the phenomenological diversity and temporal structure of LID. Recent studies have shown that specific ensembles of D1- and D2-SPNs encode specific behaviors ^15^. Consequently, different forms of LID may result from the specific activation of distinct neuronal groups including both SPN populations. To answer this question, there is a need to develop new methods enabling to monitor dyskinetic motions with high precision. In both PD patients and animal models, the classification and quantification of dyskinetic movements is based on rating scales that assign dyskinesia severity scores to different body parts during monitoring periods of 1-2 minutes. Although these rating methods are well validated for translational research ^17,18^ they do not offer the temporal resolution that would be necessary to resolve neuronal events accounting for the different motor components that make up LID.

In this study, we present a new approach to automatically quantify dyskinetic movements with high temporal resolution in freely-moving mice, which enabled us to investigate in detail the striatal pathophysiology associated with LID. We developed a semi-supervised approach using unsupervised behavioral clustering based on inertial measurement units (IMUs) and video, combined with a supervised higher-order clustering based on dyskinesia annotations. Using this approach, we were able to capture different types of dyskinesia as well as the presence of pathological rotational movements, ultimately enabling the classification of dyskinetic behavior with high accuracy. In order to investigate whether and how the activity patterns of D1-SPN and D2-SPN relate to specific dyskinesia types, we combined the usage of the IMUs with calcium imaging in freely-behaving mice. We could therefore quantify the activity patterns of D1 and D2-SPN during each dyskinesia cluster and the pathological rotations. Our results show that specific patterns of abnormal involuntary movements are encoded by specific ensembles of D1-SPNs and also D2-SPNs showing abnormally high firing activity.

## Results

### L-DOPA increases body acceleration during movement in dyskinetic mice

To render mice hemiparkinsonian, we induced chronic nigrostriatal DA depletion by injecting 6-hydroxydopamine (6-OHDA) unilaterally into the medial forebrain bundle ^9^. As expected, 6-OHDA lesioned mice exhibited a nearly complete loss of DA fibers in the ipsilateral striatum (Figure S1A, STAR Methods). All lesioned mice treated with L-DOPA (6 mg/kg/day for 3-4 days) developed axial, limb and orofacial abnormal involuntary movements (AIMs), quantified using the classical dyskinesia severity scale ^19,20^ (Figure S1B).

To study changes in motion patterns, we quantified mouse movements by measuring high-resolution acceleration and angular velocity with head-mounted wireless inertial measurement units (IMUs) and video recordings (Figure 1A, see STAR Methods). Measurements were performed in non-lesioned mice (referred to as intact, Int, n = 12) and 6-OHDA lesioned mice (Les, n = 13). Mice were placed in an open field arena, and baseline (BL) behavior was recorded for 10 min. After the BL recording, mice were injected with VEH or L-DOPA (LD) and recorded for another 10 min at 20-30 min *post* injection (timeline in Figure 1A and STAR Methods).

**Figure 1.**
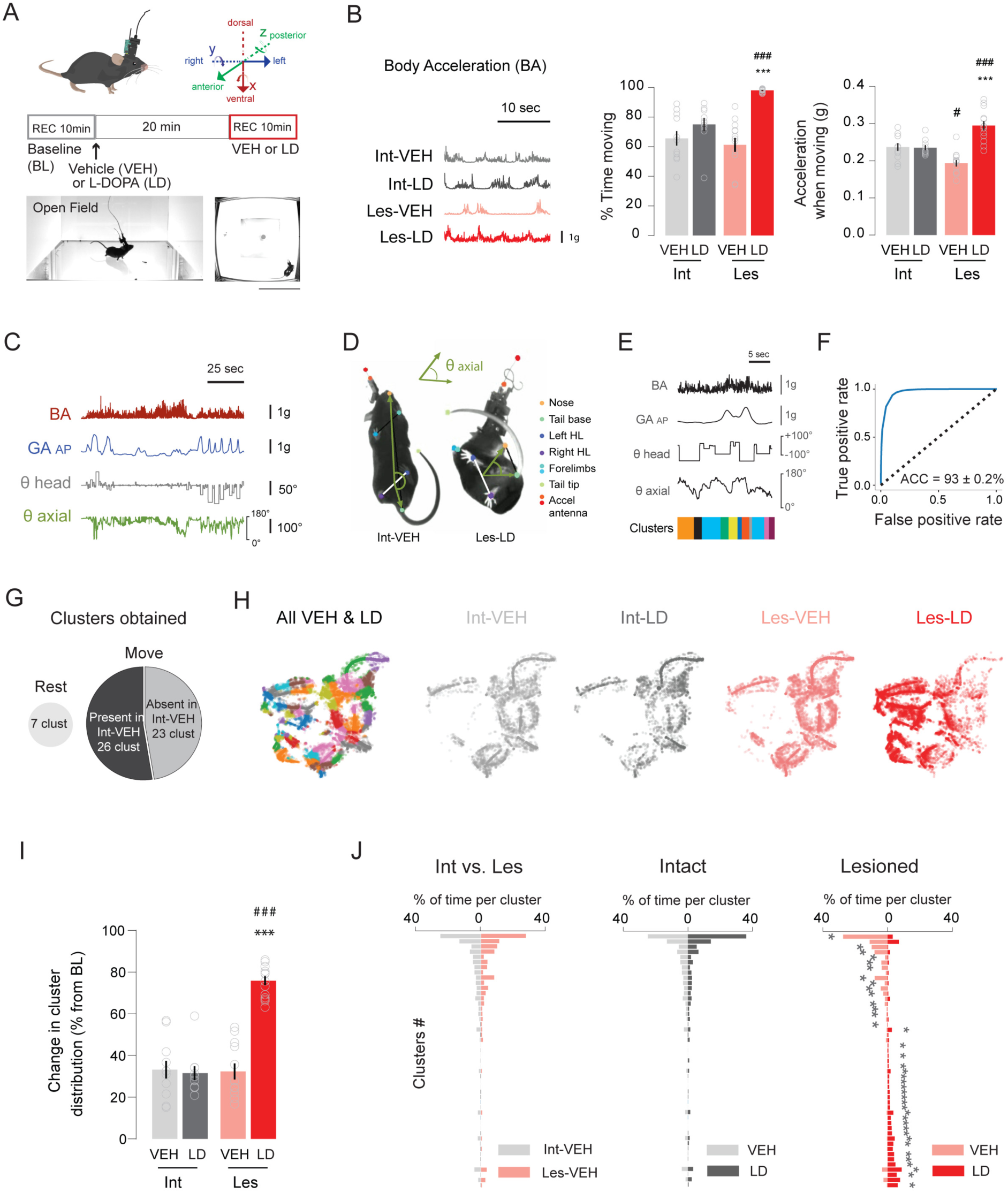
Behavioral changes in dyskinetic mice revealed by unsupervised behavioral clustering. (A) Illustration of the microendoscope and the wireless inertial measurement unit (IMU) placed on top of the mouse head. On the right is shown the orientation of the 6 axes of the IMU: three axes for the acceleration (*x* = dorso-ventral; *y* = posterior-anterior; *z* = right-left) and three axes for the gyroscope which measures the rotational velocity around these axes, represented by the rotating arrows. Below is the timeline of the experimental design (STAR Methods) and on the bottom, the view of the open field arena from the side- and bottom-placed cameras (scale bar, 20 cm). (B) Example traces of the body acceleration (BA) in the four conditions. Bar plot on the right shows the mean of the time moving as a percentage of the total time of the session ± SEM (n = 11-13 mice). Ordinary 1-way ANOVA, F_(3, 42)_ = 18.27, p < 0.001. Post hoc Bonferroni’s multiple comparisons test shows ***p < 0.001 Les-LD vs. Les-VEH; ^###^p < 0.001 Les-LD vs. Int-LD. Right: Body acceleration (g) while moving ± SEM (n = 11-13 mice). Ordinary 1-way ANOVA, F_(3, 42)_ = 17.56, p < 0.001. Post hoc Bonferroni’s multiple comparisons test shows ***p < 0.001 Les-LD vs. Les-VEH; ^###^ p < 0.001 Les-LD vs. Int-LD; ^#^ p < 0.05 Les-VEH vs. Int-VEH. (C) Example traces of the four features used for the clustering: BA, GA_AP_ (gravitational acceleration along the antero-posterior axis), θ head (head angle, proxy of rotations) and θ axial (axial bending angle, proxy of axial AIMs; see STAR Methods). (D) Video frames showing two mice (left from Int-VEH; right Les-LD) taken with the bottom camera. Shown are the 9 tracking points labeled using DeepLabCut software. The axial bending angle, θ axial, was calculated based on the hindlimbs (HL), the tail base and nose, as in ^22^ (see STAR Methods and Movie S1). Note that the Int-VEH mouse has a flat angle (∼180 deg) compared to the Les-LD mouse whose θ axial is ∼40 deg. (E) Behavioral clustering using BA, GA_AP_, θ head and θ axial. Top: BA, GA_AP_, θ head and θ axial time series. Bottom: corresponding behavioral clusters obtained by affinity propagation on similarity of discretized time series (STAR Methods). (F) Resulting clusters are well separated as indicated by a ROC analysis using a single threshold on the Earth Mover Distances (EMD). *True positive* are two segments belonging to the same cluster; *false positive* are two clusters belonging to different clusters. Original accuracy (ACC) was compared to accuracy for shuffled clusters: paired t-test, t(24) = 166.3, p < 0.001 (n = 25 mice). (G) Clustering resulted in a total of 56 clusters separated in two major groups: clusters corresponding to *rest* behavior (7 clusters) and to behavior when the animal is moving (*move,* 49 clusters). Move clusters were further subdivided into a group of clusters present in Int-VEH (26 clusters) and one absent in Int-VEH (23 clusters). (H) Two-dimensional t-SNE representation of the cluster segments. Each dot represents a behavioral segment. On the left, space representation of the VEH and LD sessions of the library (‘All VEH & LD’), each color being a different cluster. On the right, behavioral representation of the four conditions Int/VEH or LD and Les/VEH or LD. Note a very similar space representation of the segments obtained in Int-VEH, Int-LD and Les-VEH, in contrast with the ones from Les-LD. (I) Change in the cluster distributions compared to baseline (BL). Bar plot represents the percentage of change ± SEM (n = 10-13 mice). Ordinary 1-way ANOVA, F_(3, 42)_ = 43.82, p < 0.001. Post hoc Bonferroni’s multiple comparisons test shows ***p < 0.001 Les-LD vs. Les VEH; ^###^p < 0.001 Les-LD vs. Int-LD. (J) Left: percentage of time spent per cluster between Int-VEH and Les-VEH (Mann-Whitney U test for the 26 clusters present in Int-VEH; all p > 0.05 after Benjamini– Yekutieli post-hoc correction; n = 19 Int mice, n = 20 Les mice). Middle: percentage of time spent per cluster in Int-VEH versus Int-LD (Wilcoxon signed-rank test for all *moving* clusters; all p > 0.05 after Benjamini–Yekutieli post-hoc correction, n = 19 mice). Right: percentage of time spent per cluster in Les-VEH and Les-LD (Wilcoxon signed-rank test for all *moving* clusters; *p < 0.05 for 39 out of 49 clusters after Benjamini–Yekutieli post-hoc correction, n = 20 mice).

We first quantified general movement parameters such as the percentage of time the animals moved per session and their body acceleration (BA) captured with the IMUs (Figure 1B). Upon initial inspection, we observed that the open field trajectories showed different profiles between the four conditions (Figure S1C). In particular, movements were confined to one side of the arena in the Les-LD group. Although having confined movements, Les-LD mice showed a higher BA (see example traces of BA in Figure 1B). The 6-OHDA lesion did not significantly alter the percentage of time moving (Figure 1B, histogram to the left), but significantly reduced the acceleration while moving (STAR Methods) (p < 0.05 for Les-VEH vs. Int-VEH, Figure 1B, histogram to the right). The administration of L-DOPA significantly raised both the percentage of time moving and the acceleration when moving in the lesioned but not the intact animals (Figure 1B, *Acceleration when moving;* p < 0.001 for Les-LD mice vs. both Int-LD and Les-VEH). Taken together, the above data demonstrate that, while nigrostriatal DA denervation induced bradykinesia, L-DOPA induced an excessive motor output in hemiparkinsonian mice, raising both the percentage of time moving and the BA when moving. However, with the metrics used, it was not possible to extract any information about the underlying movement characteristics.

### Unsupervised clustering of motor features reveals new behavioral clusters in dyskinetic mice

To analyze the behavior in more detail and with sub-second time resolution, we used unsupervised behavioral clustering of the IMUs and video data ^15,21^. We extracted specific features from the IMU and video that capture head and body posture and motion in three dimensions. In particular, we used the following four movement features as shown in the example traces of Figure 1C: (i) total BA which distinguishes movement versus rest; (ii) gravitational acceleration along the antero-posterior axis (GA_AP_), which detects postural changes; (iii) head rotational movements along the dorso-ventral axis obtained with the gyroscope (θ head, or angular velocity); and (iv) axial bending angle extracted from the video (θ axial), which is inversely related to the degree of upper body torsion and correlates with the axial component of dyskinesia ^22^, Figure 1D, STAR Methods and Movie S1). Using a first cohort of mice (n = 12 Int, n = 13 Les, total of n = 81 sessions), we created a library of the standard repertoire of the animal behavior in our paradigm using the unsupervised clustering algorithm based on the acceleration and video data (Figure S2A, and for details of the clustering see Figure S2A-D and STAR Methods).

With the above-mentioned features, the clustering identified a total of 56 behavioral clusters (Figure 1E, G), which were well separated as shown by a ROC analysis using a single threshold to separate clusters based on the EMD (earth mover’s distance) similarity (Figure 1F). From the 56 behavioral clusters, 7 were classified as *resting,* corresponding to periods where the mice did not move, and 49 as *moving* clusters (Figure 1G, STAR Methods). From the 49 moving clusters, we found 26 clusters present in Int-VEH, which we labeled as ‘normal’ behavior, and 23 clusters absent in Int-VEH (see STAR Methods), which were considered as ‘not normal’ or pathological behavior. Since Les-LD mice did not have periods of rest (see Figure 1B), we included only the moving clusters for the further analyses to avoid the increase in time moving as a confounding factor. A two-dimensional representation (t-SNE map) of the moving clusters in space revealed a similar space representation of the clusters obtained in Int-VEH, Int-LD and Les-VEH, with a lateralization of the behavioral segments to the right of the map. This contrasted with the t-SNE map from Les-LD, which showed that segments were lateralized to the left part of the map (Figure 1H).

We then quantified changes in the distribution of the moving clusters in each group relative to their BL (Figure 1I; note that no group differences were found at BL, Figure S2E). Clusters associated with the intact condition (Int-VEH vs. Int-LD) showed a similar change vs. BL, indicating no effect of L-DOPA treatment on the moving behavior of intact mice. A very similar result was obtained from lesioned mice treated with vehicle (see Les-VEH in Figure 1I). In contrast, Les-LD animals showed a dramatic change in cluster distribution (Figure 1I, p < 0.001 vs. the other 3 groups). While these analyses indicate that clusters absent in Int-VEH (see Figure 1G) emerge specifically in Les-LD mice, they do not reveal the changes of individual behavioral clusters in the dyskinetic condition. Therefore, we next compared the percentage of time spent per cluster in intact and lesioned mice after VEH and L-DOPA administration. To this end, we added an additional cohort of n = 7 Int and n = 7 Les mice by unsupervised clustering of individual recording sessions and by matching the resulting clusters to the library (Figure S2B; see STAR Methods). As shown in Figure 1J, the lesion *per se* did not cause a significant change in the percentage of time spent per cluster (see Int-VEH vs. Les-VEH; Figure 1J, left panel). Likewise, L-DOPA did not change the percentage of time spent in each cluster in intact mice (see VEH vs. LD Intact, middle panel). In contrast, L-DOPA caused a markedly different cluster configuration in lesioned mice, with both downregulated and upregulated clusters, and even emerging ones (see VEH vs. LD Lesioned, Figure 1J, right panel). In summary, our unsupervised clustering approach revealed major changes in the behavioral structure of L-DOPA-treated mice, and the emergence of motor motifs specifically associated with dyskinetic behaviours.

### Behavioral clustering captures specific dyskinesias and pathological rotations

We next set out to determine how the changes in cluster distribution observed in Les-LD mice related to the AIMs subtypes detected and quantified with classical rating methods. To do so, we annotated two main types of AIMs well recognizable from the videos, that is, axial dyskinesia (twisted postures of the trunk toward the side contralateral to the lesion) and limb AIMs (fluttering movements of the forelimb contralateral to the lesion) ^19,20^. We annotated frame by frame the beginning and end of axial and limb AIMs (Figure 2A, Movie S2 and see STAR Methods for details). Based on the annotations, we obtained three possible combinations of dyskinesia types, i.e. *axial alone, limb alone* and *axial+limb* dyskinesia (Figure 2A).

**Figure 2.**
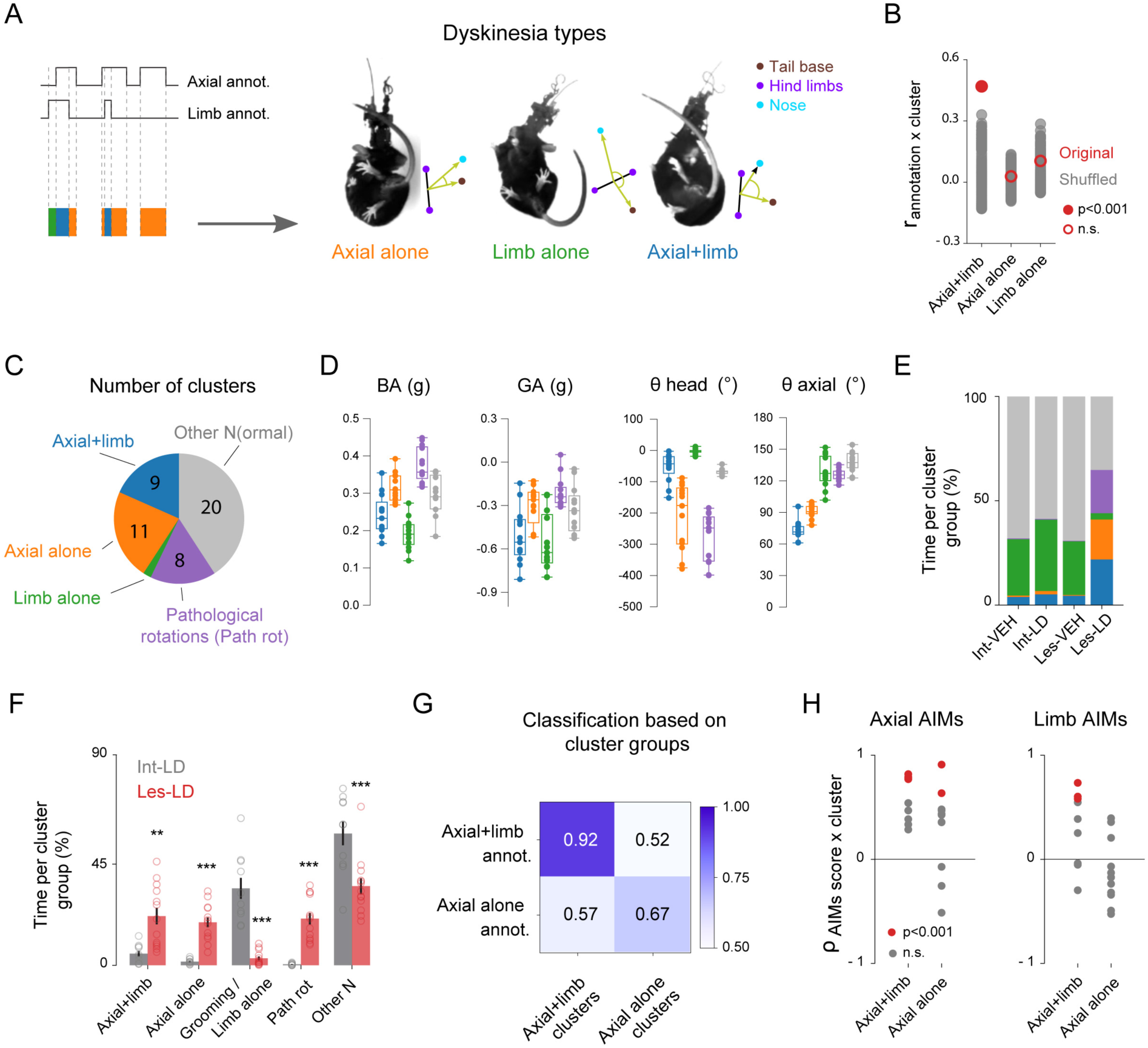
Behavioral clustering captures specific dyskinesias and other pathological behaviors. (A) Schematic representation of the frame-by-frame annotations of axial and limb dyskinesias. Three dyskinesia types were considered for the further analyses: *axial alone* (orange), *limb alone* (green) and *axial+limb* simultaneously (blue, see also Movie S1). (B) Example cluster for one mouse that significantly correlated with the *axial+limb* annotation but not with *axial alone* or *limb alone.* Bootstrap analysis of r_annotation x cluster_ to shuffled annotations (red closed circle: p < 0.001, red open circles: not significant, see STAR Methods for details). (C) Left: summary of the behavioral cluster groups obtained after correlating the behavior cluster to the dyskinesia annotations in all n = 13 mice (see STAR Methods for details). Twenty-one clusters were significantly correlated with the dyskinesia annotations: 9 were correlated with *axial+limb*; 11 to *axial alone* and 1 was correlated with *limb alone*. Twenty-eight clusters were not correlated with any dyskinesia and were denoted as *other N* for ‘normal’ (n = 20 clusters) if the clusters were present in Int-VEH mice, and *pathological rotations* (*path rot,* n = 8) if the clusters were absent in Int-VEH. (D) Feature characteristics of the cluster groups. Box and whiskers diagrams show the values of each of the 4 features (BA, GA, θ head and θ axial) corresponding to the 5 cluster groups. Values are means ± SEM (n = 13 mice). Repeated measures 1-way ANOVA, for BA, F_(2.666, 31.99)_ = 103.5, p < 0.001; for GA, F_(1.504, 18.05)_ = 21.55, p < 0.001; for θ head, F_(1.259, 15.11)_ = 80.40, p < 0.001 and for θ axial, F_(2.025, 24.30)_ = 217.4, p < 0.001. (E) Percentage of time spent per cluster group for each condition. Note that in Int-VEH, Int-LD and Les-VEH, mice spend the majority of their time (> 50%) in the *other N* cluster group and around 30% of the time in the *limb alone* cluster, which corresponds to grooming (see also Figure S2C). In Les-LD, mice show variable time spent in each of the 5 cluster groups. (F) Detailed comparison of time spent per cluster group between Int-LD (gray) and Les-LD (red). Bar plots represent the percentage of time spent per cluster group, values are the mean ± SEM (n = 10-13 mice). Two-way repeated measures ANOVA shows an effect of the *cluster groups*, F_(4,84)_ = 39.44, p < 0.001; no effect of the *lesion*, F_(1,21)_ = 0.04, p = 0.84; and an effect of the interaction, F_(4, 84)_ = 26.11, p < 0.001. Post hoc Bonferroni’s multiple comparisons test shows **p < 0.01 and ***p < 0.001 Int-LD vs. Les-LD. (G) Accuracy of a support vector classifier for predicting the type of dyskinesia (annotation) based on the *axial+limb* and *axial alone* cluster groups (see STAR Methods for details). Classification accuracies were significantly different from shuffled data. Paired t-test, *axial+limb*: t(10) = 61.1, p < 0.001; *axial alone*: t(12) = 8.8, p < 0.001. (H) Correlation between the AIM scores and the dyskinesia cluster groups *axial+limb* and *axial alone*. Each point represents a cluster. Spearman rank-order correlation, red points are clusters that are significantly correlated with the AIM scores, p < 0.001; gray points are non-significant correlations. Note that the axial AIMs are positively correlated to some of the clusters in both the *axial+limb* and *axial alone* groups. In contrast, and as expected, limb AIMs are correlated only to clusters in the *axial+limb* group but not in the *axial alone* group.

We then correlated each behavioral cluster with the three dyskinesia combinations mentioned above and grouped the clusters accordingly. We found clusters that were correlated specifically with one type of dyskinesia annotation. For example, a cluster significantly correlating with *axial+limb* annotations did not correlate with *axial alone* or *limb alone (*Figure 2B). From the 49 moving clusters, 21 were significantly correlated with the dyskinesia annotations: 9 were correlated with *axial+limb*; 11 to *axial alone* and only 1 was correlated with *limb alone* (Figure 2C and Figure S3A). Importantly, the time spent in these behavioral clusters was positively correlated to the time spent in *axial+limb* or *axial alone* annotations, but not with *limb alone* annotations (Figure S3B). The rest of the moving clusters (those not correlated to the dyskinesia annotations) included some that were present in Int-VEH mice (*other N*, 20 clusters) and others that were absent in Int-VEH, therefore considered pathological. These had a strong component of contralateral head rotations and were termed *pathological rotations* (*path rot*, 8 clusters (Figure 2C).

Importantly, the five cluster groups had composite behavioral signatures such that a single movement feature was not sufficient to differentiate between them (Figure 2D and Figure S3C, D; statistical comparisons in Table 1). For example, the two rotational components (head and axial) carried distinct information for *axial alone* and *axial+limb* dyskinesias (*cf*. blue and orange bars in Figure 2D). The *axial+limb* cluster was the most static (having the lowest BA values) and the one with strongest axial torsion (θ axial ∼70 deg, Figure 2D, Figure S3C). However, the degree of head rotation was comparable to that found in the *other N* cluster. Compared to *axial+limb*, *axial alone* was more dynamic (higher BA) and combined strong head rotations (low θ head values) with a high degree of upper body torsion (θ axial ∼90 deg). Finally, the limb cluster had low BA and GA, the lowest head rotation and low axial torsion (θ axial ∼120 deg). The *path rot* cluster group was the most dynamic of all (highest average BA, Figure 2D), while also showing the strongest head rotations (see large negative θ head values in Figure 2D). Interestingly, the turning of the head appeared to occur while the body was in a relatively straight position, as indicated by higher θ axial values compared to both “*axial alone”* and “*axial+limb*” (p < 0.001 for both comparisons, Figure 2D).

**Table 1.**
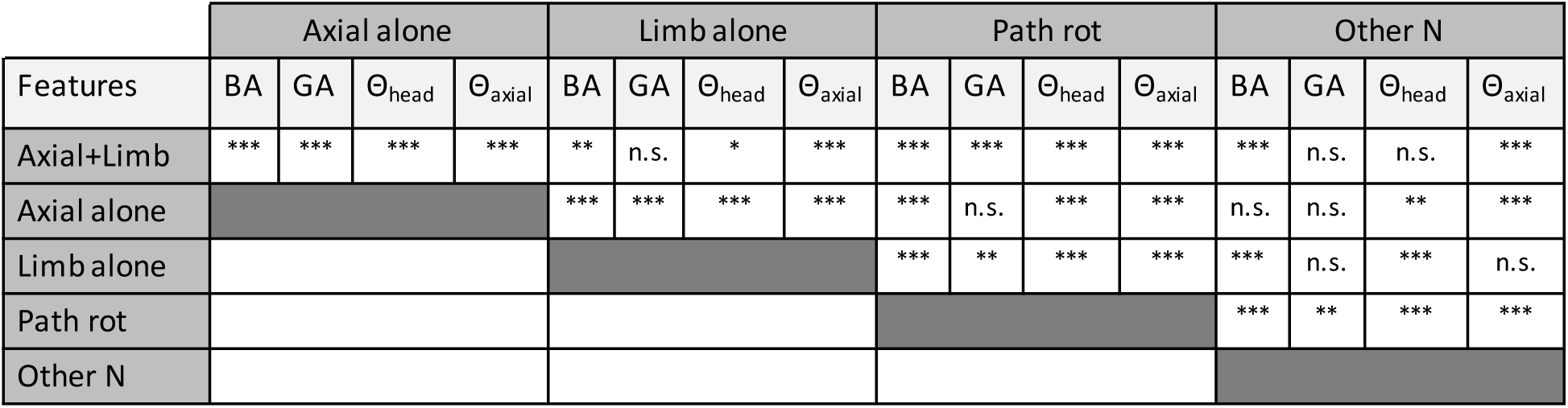
Stats related to figure 2D.

Next, we wanted to investigate how specific the dyskinesia cluster groups (*axial+limb*, *axial alone*, *limb alone*) were to the dyskinetic condition (Les-LD). We first examined the percentage of time spent per cluster group for each condition (Figure 2E). As expected, intact mice (Int-VEH and Int-LD) spent the majority of the time in the *other N* cluster group (60-70% of time during moving), and a similar result was obtained from the Les-VEH group (Figure 2E). In contrast, Les-LD mice spent the majority of the time in the dyskinesia and *path rot* cluster groups (Figure 2E). A group comparison of percentage time spent/cluster revealed that *axial+limb* and *axial alone* clusters were highly specific to the Les-LD cohort (Figure 2F). This was in stark contrast to the *limb alone* cluster, which was highly expressed also in all other cohorts. We hypothesized that limb AIMs could share kinematic features with certain phases of a grooming sequence (described by Berridge and Whishaw 1992) and therefore decided to annotate grooming in a subset of intact mice to test this hypothesis. Indeed, the *limb alone* cluster could predict grooming in intact mice with an accuracy of 73% (Figure S4A). For this reason, and because *limb alone* dyskinesia was not very prevalent in the Les-LD condition (3.1% of time during moving) and did not significantly correlate with the limb AIM scores (Figure S4B), we decided to not further analyze this behavioral cluster in the remainder of this study.

The emergence of specific dyskinesia clusters should allow one to predict different types of dyskinesias using our behavioral cluster analysis. In fact, a classifier with a “leave-one-out cross-validation” approach (see STAR Methods for details) was able to predict *axial+limb* and *axial alone* dyskinesias with an accuracy of 92% and 67%, respectively, based on the behavioral clusters (Figure 2G). While *axial+limb* and *axial alone* clusters are correlated with the occurrence of *axial+limb* and *axial alone* dyskinesias, respectively, it is not clear how these clusters relate to the classical dyskinesia scores. We therefore tested the relationship of the dyskinesia clusters to axial and limb AIMs by calculating the correlation of the time spent per individual behavioral cluster to the AIM scores during the 10 min of recording. Indeed, overall, *axial+limb* as well as *axial alone* clusters were positively correlated with the axial AIMs and a subset of clusters had a significant positive correlation (Figure 2H, red points in left panel), suggesting that some behavioral clusters only happen under more severe AIMs. Furthermore, and as expected, *axial+limb* clusters were mainly positively correlated with limb AIMs whereas *axial alone* clusters had overall negative correlations (Figure 2H, right panel). On the contrary, the correlation between *limb alone*, *path rot* or *other N* clusters and AIMs was much more variable, with the significant correlations found only in negatively correlated clusters (Figure S4B). Taken together, these results indicate that *axial+limb* and *axial alone* clusters are highly predictive of the occurrence of *axial+limb* and *axial alone* dyskinesias.

### L-DOPA oppositely modulates D1 and D2-SPNs activity in 6-OHDA lesioned but not intact mice

To determine how different dyskinetic behaviors are encoded by the striatum, we performed calcium imaging in combination with our semi-supervised behavioral clustering analysis. We used microendoscopic one-photon calcium imaging of *GCamP6f*, selectively expressed in D1-SPNs or D2-SPNs using D1-Cre and A2a-Cre transgenic mice, respectively. After 6-OHDA lesion and virus injection, mice underwent chronic GRIN (gradient index) lens implantation in the dorsolateral striatum (see extent of viral transduction, lesion and lens placement in Figure 3A and STAR Methods). After having a cleared and stable field of view of the striatum (Figure 3B), mice were placed in an open field arena and recorded for 10 min before and after VEH/LD treatment, as in *Timeline* of Figure 1A. Calcium dynamics from up to 300 neurons (Figure 3B) could be recorded simultaneously and aligned with the behavioral clusters from the semi-supervised behavior analysis (STAR Methods, and Movie S3 and S4). Intracellular calcium events per single neurons were detected using a constrained non-negative matrix factorization for microendoscopic data (CNMF-E; ^15,24,25^ see STAR Methods) and spike events were extracted using the MLspike algorithm (^26^ , STAR Methods).

**Figure 3.**
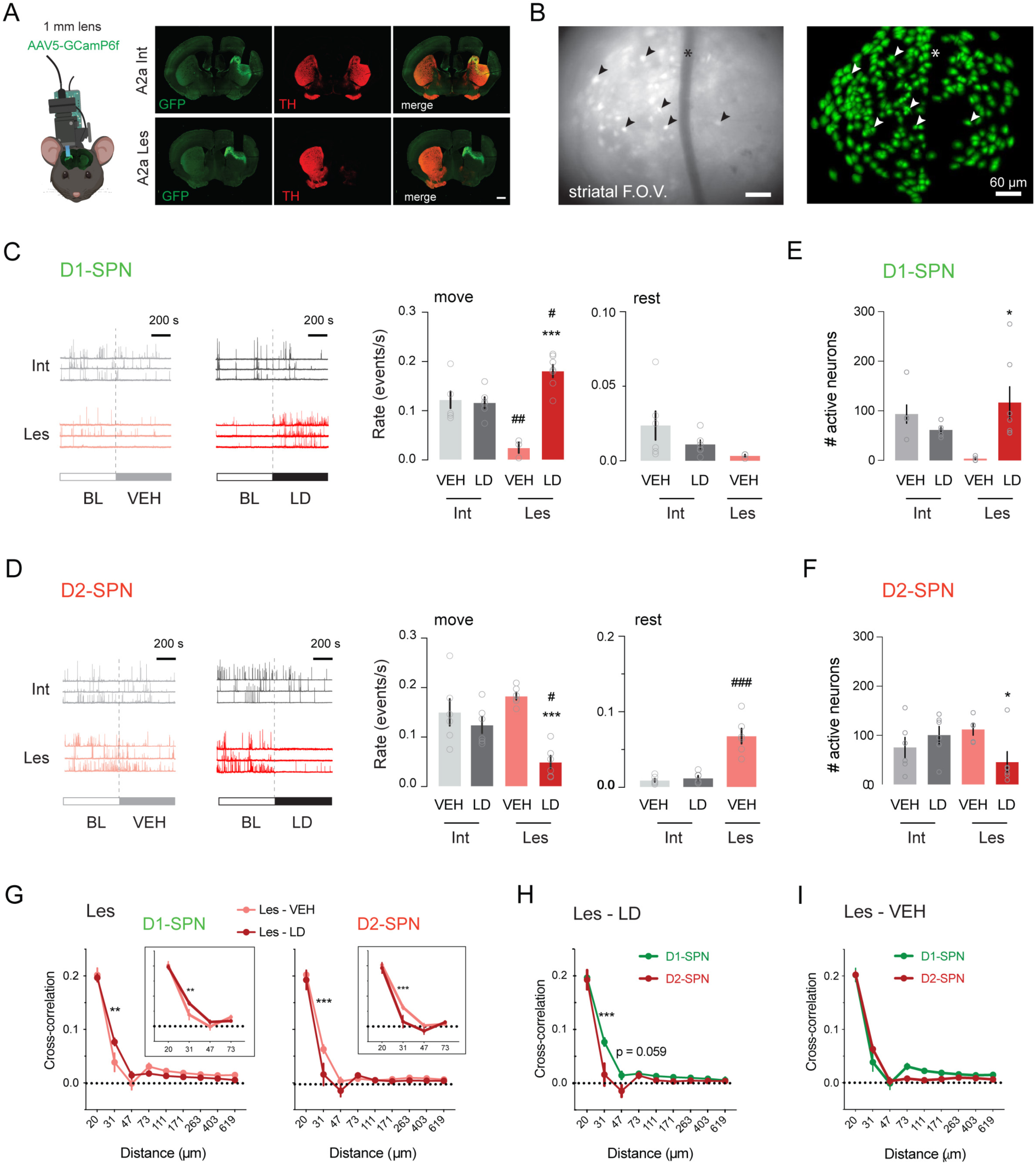
SPNs average activity is oppositely modulated by L-DOPA in 6-OHDA lesioned mice. (A) Cartoon of a mouse head showing the micro-endoscope on top of the mouse head connected to a 1 mm GRIN lens placed in the dorso-lateral striatum and the wireless IMU on the back of the micro-endoscope. On the right, coronal sections of two example mouse brains at the level of the striatum. Photomicrographs were acquired from A2a-Cre transgenic intact (A2a Int, top row) and 6-OHDA lesioned (A2a Les, bottom row) mice injected intrastriatally with the *AAV5-GCamP6f* viral vector. GFP expression (revealed with GFP antibody in green) in the dorso-lateral striatum shows the region of the striatum transduced with *AAV5-GCamP6f* viral vector, which is expressed below the 1 mm lens. TH (tyrosine hydroxylase, in red) shows the dopamine terminals in the striatum (note that A2a Les has a complete dopamine depletion shown by the lack of TH immunostaining in the right striatum). Merged photograph shows colocalization of GFP and TH showing the expression of *GCamP6f* in an intact (top) and a lesioned striatum (bottom) (scale bar: 1 mm). (B) Field of view of the striatum (striatal F.O.V.) through the lens of a D1-Cre lesioned mouse treated with LD, corresponding to a maximum projection of 3000 frames of the video recording. Fluorescent calcium signal shows increased fluorescence in neuronal somas (example neurons depicted by the arrows) and lack of fluorescence of a blood vessel (shown by an asterisk). Right picture shows the same F.O.V. with the total number of neurons (283 neurons) detected with the CNMFe algorithm during both BL and LD sessions, shown as footprints or ROIs (regions of interest) colored in green (scales: 60 µm). (C) Event rate of D1-SPNs in Int and Les mice treated with VEH and LD. On the left, example traces of n = 3 neurons of Int (gray, top row) and n = 3 neurons of Les (red, bottom row) mice (scale bar: 200 sec). First column corresponds to traces at BL and after VEH and the second column shows traces at BL and after LD (note that BL and VEH/LD are separated by a dashed line). Bar graph represents the average of the event rates (events/s) per mouse and session when moving (move) ± SEM (n = 3-7 mice per group and 6 to 18 sessions per group). Ordinary 1-way ANOVA, F_(3, 18)_ = 15.60, p < 0.001. Post hoc Bonferroni’s multiple comparisons test shows ***p < 0.001 Les-LD vs. Les-VEH; ^#^ p < 0.05 Les-LD vs. Int-LD; ^##^p < 0.005 Les-VEH vs. Int-VEH. The bar graph on the right shows the event rates when the mice are at rest. Note that the Les-LD group is not represented because Les-LD mice did not rest (see STAR Methods). Ordinary 1-way ANOVA, F_(2, 12)_ = 1.889, p = 0.1936 (see Movie S3 for D1-SPN calcium imaging aligned to the video camera recording). (D) Event rate of D2-SPNs in Int and Les mice treated with VEH and LD. On the left, example traces of n = 3 neurons of Int and Les mice as in (C). Bar graph represents the average of event rates (events/s) per mouse and session ± SEM (n = 6 mice per group and 8 to 14 sessions per group). Ordinary 1-way ANOVA, F_(3, 20)_ = 10.59, p = 0.0002. Post hoc Bonferroni’s multiple comparisons test shows ***p < 0.001 Les-LD vs. Les-VEH; ^#^p < 0.05 Les-LD vs. Int LD. Bar plot on the right shows the event rates when mice are at rest. Ordinary 1-way ANOVA, F_(2, 15)_ = 29.11, p < 0.001. Post hoc Bonferroni’s multiple comparisons test shows ^###^p < 0.001 Les-VEH vs. Int-VEH (see Movie S4 for D2-SPN calcium imaging aligned to the video camera recording). (E) Number of active (detected with CNMFe algorithm) D1-SPNs in Int and Les mice treated with VEH and LD when mice were moving. The plots show the number of active neurons ± SEM (n = 3-7 mice per group and 6 to 18 sessions per group). Ordinary 1-way ANOVA, F_(3, 19)_ = 3.561, p = 0.0338. Post hoc Bonferroni’s multiple comparisons test shows *p < 0.05 Les-LD vs. Les-VEH. (F) Number of active D2-SPNs in Int and Les mice treated with VEH and LD. The plots show the number of active neurons ± SEM (n = 6 mice per group and 8 to 14 sessions per group). Ordinary 1-way ANOVA, F_(3, 20)_ = 3.134, p = 0.0484. Post hoc Bonferroni’s multiple comparisons test shows *p < 0.05 Les-LD vs. Les-VEH. (G) Spatiotemporal cross-correlation between pairs of active SPNs of Les mice after VEH vs. LD. For D1-SPNs, two-way repeated measures ANOVA shows an effect of the *distance*, F_(8,54)_ = 195.9, p < 0.001; no effect of the *treatment*, F_(1,54)_ = 0.0088 , p = 0.9257; and an effect of the interaction, F_(8, 54)_ = 2.421, p = 0.0259. Post hoc Bonferroni’s multiple comparisons test shows **p < 0.01 at 31 µm distance between VEH and LD (squared inset from 30 to 73µm). For D2-SPNs, two-way repeated measures ANOVA shows an effect of the *distance*, F_(8,45)_ = 108.5, p < 0.001; the *treatment*, F_(1,45)_ = 8.651 , p = 0.0051; and the interaction, F_(8, 45)_ = 2.750, p = 0.0146. Post hoc Bonferroni’s multiple comparisons test shows ***p < 0.001 at 31 µm distance between VEH and LD (squared inset from 30 to 73µm). (H) Comparison of the spatiotemporal cross-correlation between pairs of active D1-SPNs vs. D2-SPNs of Les mice after LD. Two-way repeated measures ANOVA shows an effect of the *distance*, F_(8,88)_ = 193.8, p < 0.001; a smaller effect of the *treatment*, F_(1,11)_ = 4.984 , p = 0.0473; and an effect of the interaction, F_(8,88)_ = 4.741, p < 0.001. Post hoc Bonferroni’s multiple comparisons test shows ***p < 0.001 at 31 µm distance between D1-SPN and D2-SPN. (I) Comparison of the spatiotemporal cross-correlation between pairs of active D1-SPNs vs. D2-SPNs of Les mice after VEH. Two-way repeated measures ANOVA shows an effect of the *distance*, F_(8,88)_ = 246.5, p < 0.0001; no effect of the *treatment*, F_(1,11)_ = 0.4845 , p = 0.5008; and an effect of the interaction, F_(8,88)_ = 2.92, p < 0.01. Post hoc Bonferroni’s multiple comparisons test shows no significant difference at any distance between D1-SPN and D2-SPN.

We first sought to investigate how D1-SPNs and D2-SPNs average calcium activity was modulated by L-DOPA in lesioned and intact mice, when the mice were moving or resting (STAR Methods). When comparing the D1-SPN event rate between groups during the moving periods, we found a significant decrease in the Les-VEH group (p < 0.01 vs. Int-VEH) and a significant increase after L-DOPA in lesioned but not intact mice (see example traces of D1-SPN calcium signal and the corresponding event rates in Figure 3C). We then quantified the SPN activity during periods of rest. We could not quantify the event rate in the Les-LD condition, since dyskinetic mice did not sufficiently rest, being more than 98% of the time per session moving, as shown in Figure 1B. We found no significant changes in D1-SPNs in intact mice after L-DOPA when mice were at rest, with values similar to lesioned mice treated with VEH (Figure 3C). Next, we quantified the calcium event rate in D2-SPNs during movement, finding no effect of L-DOPA in intact mice as opposed to a significant decrease in lesioned mice (see example traces and quantification in Figure 3D; see also the ratio D1-SPN/D2-SPN activity in Figure S5A, and a summary Table in Figure S5B). Interestingly, the D2-SPN activity during resting periods was very low in intact mice (both the Int-VEH and Int-LD cohorts), but markedly high in the Les-VEH group (p < 0.001 for Les-VEH vs. Int-VEH, Figure 3D). We also quantified the relative change in event rate from BL and captured the opposite and significant modulation of SPNs by L-DOPA in lesioned mice when moving (Figure S5C). No differences in the relative change in event rate were found in resting periods between Int-VEH vs. Int-LD, and similar change was observed in Les-VEH. Overall, these results demonstrate that L-DOPA oppositely modulates the average activity of both SPN populations during movement in lesioned mice, increasing the calcium event rate in D1-SPN while decreasing it in D2-SPNs. In addition, also the number of active neurons was oppositely modulated by L-DOPA in the two SPN populations. Thus, the number of active D1-SPNs increased while the number of active D2-SPNs decreased in lesioned mice treated with L-DOPA (Figure 3E-F, p < 0.05 for Les-LD vs. Les-VEH in both SPN populations; see the changes relative to BL in Figure S5D, and the D1-SPN/D2-SPN active neuron ratio in Figure S5E), in line with a recent study from Maltese and colleagues ^27^.

Finally, we quantified the spatial cross-correlation of calcium events between pairs of active neurons as in ^15^. In all experimental groups, the cross-correlation between active SPN pairs declined sharply between 20 and 47 µm of intercellular distance (Figure 3G-I, S5F-H). In both types of SPN, no significant difference in spatial cross-correlation was found when comparing Int-VEH and Les-VEH (data not shown), indicating that the lesion *per se* did not affect the coactivity index in either SPN population. However, when examining the effect of L-DOPA, we found that pairs of D1-SPNs significantly increased their coactivity while pairs of D2-SPNs significantly decreased their coactivity specifically at ∼ 30 µm of intercellular distance (Figure 3G), and only in the lesioned cohort (see data from intact mice in Figure S5F). When comparing the two SPN categories in the Les-LD group, a large difference in cross-correlation emerged at the same critical distance, where the coactivity of D1-SPN pairs was approximately 5-fold larger than that of D2-SPN pairs (Figure 3H). No significant differences between SPN categories were found after VEH (Figure 3I), nor in intact mice (Figure S5G and H). These findings suggest that the spatiotemporal activation pattern of nearby D1 vs. D2 SPN ensembles changed substantially during LID, but only at intercellular distances lower than 40 µm.

### Specific SPN ensembles are hyperactive during dyskinesia

The above analyses confirm the expected opposite changes in SPN neuronal activity observed in lesioned mice developing dyskinesia on L-DOPA, as shown in previous studies ^4–6^. However, the pattern of D1-vs. D2-SPN activity underlying specific dyskinetic behaviors has thus far remained unexplored. We therefore set out to determine how D1-SPNs and D2-SPNs encoded the behavioral clusters of dyskinetic mice. As a first approach, we focused on the Les-LD condition and calculated the average activity of both SPN populations in each behavioral cluster group, i.e. *axial+limb*, *axial alone*, *path rot* and *other N*. We found no significant differences in the average event rate of D1-SPNs or D2-SPNs between the different cluster groups (Figure 4A). In all behavioral clusters, the ratio of event rates between D1-SPNs and D2-SPNs was 1.5 to 3.5-fold larger than the average values measured in Int-LD mice when moving (see hatched line in Figure 4A), with the largest increase seen in the *axial alone* and *path rot clusters* (Figure 4A right panel). This data suggests that overall increases and decreases in D1-SPN and D2-SPN activities induced by L-DOPA are not driven by specific behavioral cluster groups.

**Figure 4.**
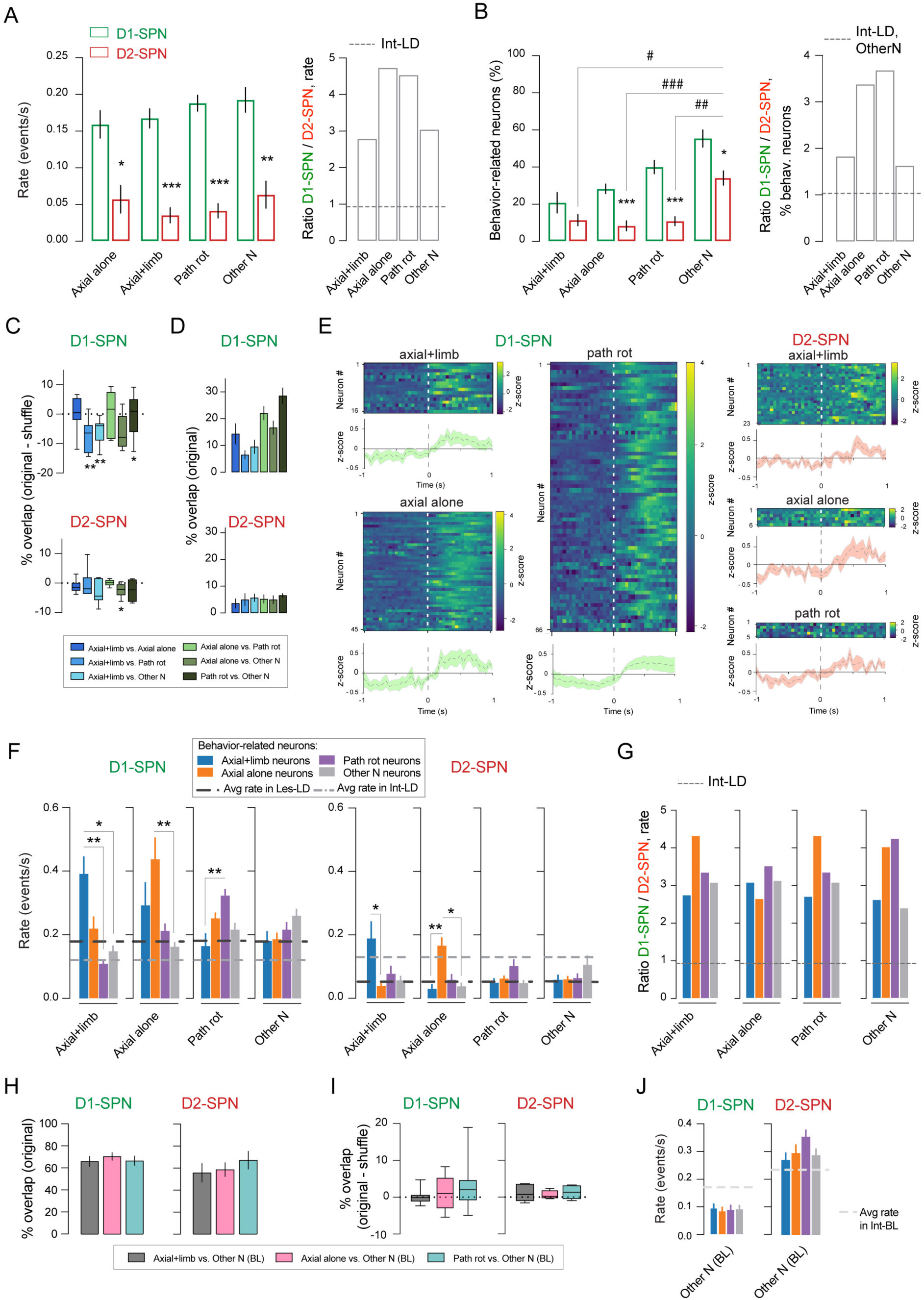
Specific sets of D1-SPNs and D2-SPNs are associated with the dyskinesia clusters. (A) Left: average event rate of all D1-SPNs and all D2-SPNs during each behavioral cluster group in Les-LD mice. The bar plots represent the mean ± SEM (n = 6-7 mice, 2-3 sessions per mouse) of D1-SPNs (green) and D2-SPNs (red) event rates (events/s) in *axial+limb*, *axial alone*, *path rot*, *other N* cluster groups. Two-way repeated measures ANOVA: *SPN type*, F_(1, 11)_ = 49.31, p < 0.001; *Cluster group*, F_(2.154, 23.69)_ = 3.004 , p = 0.0654; interaction, F_(3, 33)_ = 2.021, p = 0.13. Post hoc Bonferroni’s multiple comparisons test shows *p < 0.05 D1-SPN vs. D2-SPN in *axial+limb* cluster group; ***p < 0.001 D1-SPN vs. D2-SPN in *axial alone* and *path rot* cluster group, and **p < 0.01 D1-SPN vs. D2-SPN in *other N*. Right: ratio of the event rate in D1-SPN vs. D2-SPN per cluster group. Dashed line is the ratio between D1-SPN and D2-SPN average rate in Int-LD during move. (B) Left: percentage of behavior-related D1-SPNs and D2-SPNs in Les-LD mice (a neuron is defined as behavior-related if it showed a significant positive correlation between its activity and the behavior, see STAR Methods). The bar plots represent the mean ± SEM (n = 6-7 mice, 2-3 sessions per mouse) of the percentage of neurons significantly modulated in *axial+limb*, *axial alone*, *path rot*, *and other N* cluster groups. Dashed line is the ratio between D1-SPN and D2-SPN’s % of behavior-related neurons in Int-LD during *other N* cluster groups. Two-way repeated measures ANOVA: *SPN type*, F_(1, 11)_ = 61.44, p < 0.001; *Cluster group*, F_(1.650, 18.15)_ = 23.20 , p < 0.001; interaction, F_(3, 33)_ = 2.253, p = 0.1005. Post hoc Bonferroni’s multiple comparisons test shows: ***p < 0.001 D1-SPN vs. D2-SPN in *axial alone* and *path rot* cluster group; *p < 0.05 D1-SPN vs. D2-SPN in *other N* cluster group; in D2-SPN ^#^p < 0.05 *axial+limb* vs. *other N*; ^###^p < 0.001 *axial alone* vs. *other N*; ^##^p < 0.01 *path rot* vs. *other N*. Right: ratio of the behavior-related D1-SPN over D2-SPN per cluster group. Dashed line represents no changes in the ratio. (C) Percentage of shuffle-corrected overlap between behavior-related neuronal groups for D1-SPNs and D2-SPNs. The overlap was calculated as the number of neurons shared over the total number of neurons in the two respective behavioral cluster groups. This number varies between 0 (no shared neuron) and 100% (complete overlap). Shuffle corrected overlap is the overlap minus the shuffle overlap (see STAR Methods for details). Box and whiskers diagrams show the values for the overlap between all the different comparisons. Individual unpaired t-tests showed a significant difference (compared to 0), **p < 0.01: in *axial+limb* vs *path rot* and *axial+limb* vs *other N*; *p < 0.05: in *path rot vs other N*. (D) Percentage of overlap between the different behavioral clusters in the original data (i.e., without shuffle correction). The bar plots represent the mean ± SEM (n = 6-7 mice, 2-3 sessions per mouse). (E) Example heatmaps of individual D1-SPN (left) and D2-SPNs (right) aligned to the beginning of the behavioral events (at time point 0 s) for *axial+limb*, *axial alone* and *path rot* behavioral clusters, and the corresponding average curves of the z-scored DF/F. Unpaired t-test comparing time period (-1 to -0.1 sec) and (0.1 to 1 sec) shows significant difference in *axial+limb*, *axial alone* and *path rot* in D1-SPNs (*p < 0.05) and D2-SPNs (**p < 0.01). (F) Behavior-related D1-SPN and D2-SPN event rate during the different behavioral cluster groups. Each bar represents the average event rate ± SEM (n = 6-7 mice, 2-3 sessions per mouse) of the behavior-related neurons for each behavioral cluster group. Black dashed line is the average rate in Les-LD during move, and the gray dashed line is the average rate in Int-LD during move. Note that the behavior-related D1-SPNs and D2-SPNs show the highest activity during their respective behavior. Kruskal-Wallis nonparametric test was performed for each of the 4 cluster groups individually. Left: for D1-SPN, Kruskal-Wallis test showed significance for *axial+limb* (**p < 0.01), *axial alone* (**p < 0.01) and *path rot* (*p < 0.05) cluster groups, but not for *other N*. Post hoc Dunn’s multiple comparison shows: in *axial+limb* cluster group, **p < 0.01 ‘axial+limb neurons’ vs. ‘path rot neurons’ and *p < 0.05 vs. ‘other N neurons’; in *axial alone* cluster group, **p < 0.01 ‘axial alone neurons’ vs. ‘other N neurons’; and in *path rot* cluster group, **p < 0.01 ‘path rot neurons’ vs. ‘axial+limb neurons’. Right: for D2-SPN, Kruskal-Wallis test showed significance for *axial alone* cluster group (**p < 0.01), but not for *axial+limb* (p = 0.05, n.s.), *path rot or other N*. Post hoc Dunn’s multiple comparison shows: in *axial+limb* cluster group, *p < 0.05 ‘axial+limb neurons’ vs. ‘axial alone neurons’; in *axial alone* cluster group, **p < 0.01 ‘axial alone neurons’ vs. ‘axial+limb neurons’ and *p < 0.05 vs. *other N* cluster group. (G) Ratio of the event rate in D1-SPN vs. D2-SPN per cluster group. Dashed line is the ratio between D1-SPN and D2-SPN average rate in Int-LD during move. (H) Percentage of overlap between *axial+limb*, *axial alone*, *path rot* SPNs under LD and *other N* SPNs during BL, using the original data, i.e.,without shuffle correction (shown are the mean ± SEM; n = 6-7 mice, 2-3 sessions per mouse). (I) Percentage of overlap between the neuron groups in (H) with shuffle correction, calculated as in (C). Box and whiskers diagrams show the values for the overlap between all the different comparisons. Individual unpaired t-tests showed no significant difference (compared to 0) in any of the compared groups. (J) Behavior-related D1-SPNs and D2-SPNs event rate during *other N during BL* average event rate ± SEM; (n = 6-7 mice, 2-3 sessions per mouse). Dashed line is the average rate in Int-BL for D1-SPN and D2-SPN.

Although no differences were found in the average event rate of both SPNs between behavioral clusters, we observed that specific sets of D1-SPNs and D2-SPNs showed a significant positive correlation between their activity and the behavioral clusters (i.e., their activity increases during the corresponding behavior cluster). We refer to those neurons as ‘behavior-related neurons’ (Figure 4B). In our field of view we detected an average of 164 D1-SPNs and 176 iSPNs (all SPNs detected during both BL and LD sessions). In the D1-SPN population, 21% of neurons were positively correlated to *axial+limb*, 28% to *axial alone*, 40% to *path rot*, and 55% to *other N* cluster groups during the LD condition. In contrast, the percentage of behavior-related D2-SPNs was much smaller, with approximately 11, 8, 11 and 34% of D2-SPNs being positively correlated to *axial+limb*, *axial alone*, *path rot* and *other N*, respectively. Interestingly, the percentage of D2-SPNs associated with the *other N cluster* was at least 3-fold larger than those associated with *axial+limb*, *axial alone* and *path rot* (p < 0.05 in all these comparisons, Figure 4B). This data points to a larger recruitment of D2-SPNs during the expression of relatively normal motions despite the ongoing effect of L-DOPA. The ratio between behavior-related neurons in the D1-SPN and D2-SPN populations was strongly biased towards D1-SPNs in each cluster, with the largest increase relative to Int-LD values (> 2-fold) in *axial alone* and *path rot* (Figure 4B right panel).

*Axial+limb* and *axial alone* dyskinesia share some behavioral characteristics (i.e., the axial component) that could be reflected in an overlap between the neuronal ensembles related to the two types of dyskinesia (i.e., *axial+limb* vs. *axial alone*). We, therefore, investigated the percentage of overlap between the two cluster groups, corrected by the overlap in shuffled data that would be expected purely by chance if there was no common set of neurons between the *axial+limb* and *axial alone* neuronal groups. In addition, we compared the overlap between the neuronal groups associated with the other behaviors observed during LD (Figure 4C). This analysis revealed that the overlap between any of the behavior-related D1-SPNs and D2-SPNs during LD is not more than the one expected by chance (i.e., no significant positive increase from 0; 0% overlap corresponding to no shared neurons and 100% to a complete overlap between the neuronal groups). In D1-SPNs, we found significant negative values, indicating that between three behavior-related neuronal groups (*axial+limb* vs. *path rot*, *axial+limb* vs. *other N* and *axial alone* vs. *other N*) there is less overlap than would be expected by chance. We calculated the percentage of this overlap and found that the overlap across behavioral clusters ranged from 6 to 27% for D1-SPNs and from 4 to 7% for D2-SPNs. This data shows that the overlap between active neurons across behavioral clusters is not different from chance level, therefore indicating that the behavior-related neurons are specific to their cluster group. Accordingly, the activity of specific SPN ensembles was strongly modulated at the onset of dyskinetic and *path rot* behavioral events, shown by the example heatmaps of the individual SPN activity aligned to the start of the behavioral event and the corresponding average curves (Figure 4E). We found a significant difference between the time period before (-1 to -0.1 sec) and after (0.1 to 1 sec) the start of *axial+limb*, *axial alone* and *path rot* in both D1-SPNs and D2-SPNs.

We next compared calcium event rates in these specific sets of SPNs in Les-LD mice (Figure 4F). The behavior-related D1-SPNs showed an abnormally high activity in all clusters when compared to the average rate in Int-LD or Les-LD (see dashed lines in Figure 4F). The largest increase occurred in D1-SPNs tuned with the *axial+limb* and *axial alone* clusters (see blue and orange bars in Figure 4F, left). Compared to the Les-LD condition (dark dashed line), behavior-related D2-SPNs also showed a significant increase in event rate specifically during their associated cluster (Figure 4F right; 4G shows the ratio between D1-SPN and D2-SPN’s rate), in some cases exceeding the average values measured in intact mice (see *axial+limb* and *axial alone* clusters in Figure 4F right). Our data indicate that, while cluster-related SPNs fire also during other behaviors, they show their maximal event rate during their specific behavioral cluster. Overall, these results suggest that each type of dyskinetic behavior is encoded by a specific subset of D1-SPNs firing at a frequency at least 2-fold larger than the average D1-SPN activity rate in the Les-LD condition (∼0.4 Hz), along with a smaller subset of D2-SPNs showing an approximately 3-fold increase above the average D2-SPN activity in lesioned animals, yet firing at a lower rate than the coactivated D1-SPNs (∼0.18 Hz).

Previous studies have identified dyskinesia-specific D1-SPNs that are exclusively activated during LID ^4^. To gain better insight into the emergence of dyskinesia-specific ensembles, we took advantage of our experimental paradigm that allowed for extracting the activity of dyskinesia-specific SPNs also during normal behavior. Under BL condition, about 60% of SPNs were associated with clusters of motor features emerging during normal movement (data not shown), in line with previous findings from intact mice ^15^. Next, we compared the overlap between the *other N (BL)* neurons and dyskinesia-specific neurons in Les-LD (Figure 4H, where *other N (BL)* corresponds to *other N* neurons during BL condition). We found a high overlap between both the dyskinesia and *path rot* neuronal groups with the SPNs encoding for *other N (BL)* clusters, amounting to 61% and 68% overlap on average for D1-SPNs and D2-SPNs, respectively (Figure 4H). Although this overlap was statistically not different from shuffled data (Figure 4I), these results indicate that the majority of SPNs associated with dyskinesia and *path rot* encode for normal behaviors during non-dyskinetic periods. We finally measured the neural activity of *axial+limb*, *axial alone* and *path rot* neurons during BL, and found a similar event rate, comparable to that measured under BL conditions in intact mice (Figure 4J).

The finding that the dyskinesia and *path rot* neurons were indeed active during *other N* behavioral clusters at BL, raised the question of whether *other N* behavioral clusters during which these neurons were active would show any behavioral similarity to the dyskinesia or *path rot* clusters. To address this question, we compared the EMD similarity of behavioural clusters recorded in the Les-LD condition vs. BL clusters during which the same neurons were active, considering *axial+limb*, *axial alone,* and *path rot* SPNs (Figure S6A). Specifically, for each SPN group, we identified the dyskinesia/*path rot* clusters to which they were tuned (Les-LD) and compared their EMD similarity to *other N* clusters (Les-BL) during which the same neurons were positively modulated (“pos mod”) versus the remaining clusters in the *other N* group (“other”). Interestingly, we observed that the EMD similarity between the *axial+limb* clusters and the “pos mod” clusters was significantly higher for both SPNs than the one between the *axial+limb* clusters and the “other” clusters in the *other N* category (Figure S6A). However, this significant difference was not observed in *axial alone* and *path rot*. These results indicate that the behaviors encoded by the *axial+limb* neurons during BL are more similar to *axial+limb* dyskinesia than the ones encoded by other neurons. In turn, this suggests that the neurons showing largest activation during a given type of dyskinetic motion encode for related forms of body movements during normal behaviors.

## Discussion

We here present a new approach to automatically detect dyskinetic movements with sub-second resolution in freely-moving mice. We developed a semi-supervised approach using unsupervised behavioral clustering based on inertial measurement units and video, combined with a supervised group clustering based on dyskinesia annotations. Using this approach, we were able to capture different types of dyskinesia and other behaviors absent in intact mice classified as pathological rotations. In order to investigate the interplay between D1-SPNs and D2-SPNs during specific dyskinesia types, we combined the new behavioral method with SPN calcium imaging in freely-behaving mice. We could therefore quantify the activity patterns of D1-SPN and D2-SPN during each dyskinesia cluster. Our results show that the two dyskinesia types and the pathological rotations were encoded by highly specific sets of D1-SPNs and D2-SPNs. These sets of SPNs emerge from combinations of D1 and D2-SPNs encoding normal behaviors under baseline conditions.

The present study introduces the first application of a non-invasive miniaturized wireless IMU version for mice to quantify pathological movements. The unsupervised behavioral clustering method was based on previous work developed by Klaus and collaborators ^15^, where the body movement of healthy mice in an open field arena was monitored and clustered using wired IMUs and video data. As in Klaus and colleagues, we used total BA and GA in the antero-posterior axis; however, diverging from their approach, the head rotation parameter was taken from the IMUs and in particular from the gyroscope component, rather than from the video. The addition of the fourth feature, the axial bending angle, was key to capture the axial component of dyskinesia ^22^. Using the combination of these four features, the unsupervised behavioral clustering detected significant changes in the cluster distribution of dyskinetic mice, evidenced by the emergence of a new behavioral space represented in Figure 1H. Our approach enabled us to prove that LID is not simply an acceleration or exaggeration of normal movements, but involves the emergence of abnormal motor motifs that are not found in non-dyskinetic animals, whether intact or parkinsonian.

To establish the nature of the obtained behavioral clusters, we used the dyskinesia annotations to group the clusters into higher order groups. As a result, we obtained cluster groups corresponding to three specific dyskinesia types, i.e. axial combined with limb, axial alone and limb alone. The latter was discarded from further analyses due to a large degree of overlap with the kinematic properties of limb movements during grooming in intact animals. Based on the combined use of four relevant primary features, the behavioral signatures identified by the cluster groups represent with high fidelity the posture-motion dynamics of dyskinetic mice exhibiting axial and limb AIMs. For example, as dyskinetic mice develop limb AIMs, they stop locomoting to execute the characteristic fluttering movement of the contralateral forelimb. This was reflected in the *axial+limb* cluster group by a decrease in the BA and rotations, along with a position of the head closer to the floor (lower GA). When dyskinetic mice develop axial AIMs without limb involvement, we would expect an increase in BA and head rotation combined with a twisted, dystonic body posture (low axial angle) and with the head raised towards a higher GA level. These features were indeed observed in the *axial alone* clusters. In addition, the cluster expression of *axial-limb* and *axial alone* was significantly correlated with the corresponding AIMs scores. These results show that our behavioral clustering method can detect and quantify dyskinetic motor patterns with very high kinematic precision.

Interestingly, our clustering analysis detected an abnormal behavior that we defined as pathological rotations (*path rot*). The *path rot* cluster group included behavioral motifs that were not correlated to classical axial and/or limb AIMs, yet pathological because totally absent in intact and Les-VEH mice. This cluster group was characterized by pronounced head rotation (large head angle) occurring simultaneously with high body acceleration. The axial bending angle was however comparable to that measured in the *other N* cluster group. *Path rot* behavior most likely corresponds to an initial phase of axial dyskinesia, which typically starts with a tight contralateral twisting of the head preceding the torsion of the body and the stopping of forward body motions. Contralateral rotations are frequently measured in unilateral models of PD-LID using automated videotracking or rotary sensors with photobeam detectors. Measured in this way, contralateral rotations correlate poorly with LID ratings ^28^, and they are indeed induced to a much larger degree by non-dyskinesiogenic treatments with DA agonists compared to L-DOPA ^28,29^. In many circumstances, contralateral rotations have been found to reflect therapeutic-like effects of antiparkinsonian or antidyskinetic treatments (discussed in Cenci and Crossman 2018). Importantly, the high kinematic precision offered by our method enabled us to isolate pathological components of the animal’s rotational behavior, where an abnormally tight turning of the head occurs while the mouse is moving forward (high body acceleration).

In the second part of our study, we aimed at revealing the neural activity patterns of D1-SPN and D2-SPN during dyskinesia. As previously reported ^5,6^, dyskinetic mice exhibited an increased average D1-SPN activity and a reduced D2-SPN activity during periods of movement. We also investigated whether SPNs would exhibit altered spatiotemporal activation patterns, as indicated by Parker and colleagues ^5^. Overall, cross-correlations between pairs of active SPNs exhibited the same spatiotemporal configuration in all experimental groups, with a sharp decline between 20 and 47 µm and hardly any cross-correlation at intercellular distances larger than 100 µm. This suggests that small groups of contiguous SPN fire together also under parkinsonian and dyskinetic conditions, presumably driven by the same excitatory input ^30^, resembling the situation reported in intact mice during movement ^15,31^. At variance with the results of ^5^, our data therefore indicate that the gross spatial organization of SPN activity is maintained during LID, although the number of active neurons, their event rates, and the D1-SPN/D2-SPN activity ratio are profoundly abnormal. Importantly, however, a marked difference between SPN types was found specifically in the Les-LD mice at intercellular distances below 40 µm, with a larger co-activation of D1-SPNs compared to D2-SPNs. Given that co-activity patterns of D1 and D2-SPNs were basically identical under normal conditions (see Figure S5G-H), the highly significant divergence in cross-correlation between the two SPN types at short intercellular distances represents a substantial change. This data points to an altered interplay between contiguous groups of D1- and D2-SPNs that should be equally coactive during movement ^15^.

For the first time, we could relate specific dyskinetic movements to the neural activity of striatal ensembles with single-cell resolution. At first, we saw that average calcium event rates of D1-SPNs and D2-SPNs did not differ significantly between behavioral clusters. This data indicated that the average increase in D1-SPN and decrease in D2-SPN activity was not driven by a specific behavior, but also that the average SPN activity *per se* could not explain the emergence of different types of dyskinesia. Next, taking advantage of the single-cell resolution, we could identify specific non-overlapping groups of D1-SPNs and D2-SPNs whose activity was correlated to each behavioral cluster group. Although these specific sets of neurons fired also under other conditions, their activity was maximal during their associated behaviors. The sets of D1-SPNs linked with the two dyskinesia cluster groups had doubled their activity compared to the average activity during the moving periods. Although the entire D2-SPNs population showed very low calcium event rates in Les-LD, the D2-SPN ensembles associated with dyskinesia or pathological rotation exhibited markedly higher activity than the rest of the D2 population during the expression of their corresponding behavioral clusters. This was particularly clear for D2 SPNs tuned to the *axial+limb* and *axial alone* clusters, which showed a 3-fold increase in event rate compared to the average value found in Les-LD (and also higher than the Int-LD average rate) during the expression of those specific behaviors. However, despite showing a relatively high activity, these D2-SPN groups fired at an approximately 2-fold lower rate than the D1-SPNs tuned to the same behavioral cluster. The marked hyperactivity of certain groups of D1-SPN over the average of that population is likely to depend on a stronger excitatory drive as shown in ^30^. Importantly, we found that the majority of the SPNs associated with the dyskinesia and pathological rotation clusters were also active during normal behaviors at baseline, suggesting that their marked activity levels during LID might trigger a shift from normal to abnormal motion patterns. Under baseline conditions, these neurons appear to encode for behaviors that are physically similar to the type of dyskinesia to which they are tuned, suggesting that the predetermined phenotype of hyperactive neurons determines which pattern of dyskinesia will emerge on L-DOPA.

In summary, using a novel approach to quantify dyskinetic movements, our study unveils the underlying changes in D1-SPN and D2-SPN activity with unprecedented granularity. Using in vivo recordings with single-cell resolution, we not only confirm the existence of specific sets of D1-SPNs associated with dyskinetic motions, but also describe the presence of a small group of D2-SPNs that are disproportionately active in each dyskinesia cluster. The large disparities in event rates and cross-correlation found between D1- and D2-SPNs indicate that, despite their relatively high activity levels, dyskinesia-specific D2-SPNs fail to brake nearby hyperactive D1-SPNs via inhibitory axon collaterals. While confirming that an imbalance between D1-vs. D2-SPN activity in favor of the former is a generic signature of LID, our results indicate that specific combinations of abnormally active SPNs dictate the moment-to-moment expression of particular LID features. Interestingly, the SPN ensembles tuned to specific dyskinetic motions appear to encode physically related normal movements under baseline conditions.

## METHODS

### EXPERIMENTAL MODEL AND SUBJECT DETAILS

The study was performed in bacterial artificial chromosome (BAC) transgenic mice expressing Cre recombinase under the control of the dopamine D1 receptor (D1-Cre, Tg(Drd1a-cre) FK150Gsat/Mmucd; MMRRC #029178-UCD) for targeting of direct-pathway SPNs, and adenosine A2a receptor (A2a-Cre, B6.FVB(Cg)-Tg(Adora2acre) KG139Gsat/Mmucd; MMRRC #036158-UCD) for targeting indirect-pathway SPNs. All lines have been backcrossed onto C57Bl6/J mice for at least 8 generations. Experimental mice were 3 to 5-month-old males housed on a 12-hr light/dark cycle with *ad libitum* access to food and water. All animal procedures were reviewed and performed in accordance with the Champalimaud Center for the Unknown Ethics committee guidelines and approved by the Portuguese Veterinary General Board (Direçao Geral de Veterinária, Ref. No. 0421/000/000/2014). Sample size is detailed in the Results or figure legends.

### METHOD DETAILS

#### 6-Hydroxydopamine lesion and virus injection

6-hydroxydopamine (6-OHDA) lesion and virus injection were performed in the same surgery. Surgeries were performed under isoflurane (1%-3%, plus oxygen at 1-1.5 l/min) anaesthesia on a stereotactic frame (David Kopf Instruments, Model 962LS), with a mouse adaptor (David Kopf Instruments, Model 923-B Mouse Gas Anaesthesia Head Holder). Throughout each surgery, mouse body temperature was maintained at 37°C using an animal temperature controller (ATC1000, World Precision Instruments) and afterwards, each mouse was allowed to recover from the anesthesia on a heating pad. After shaving and disinfecting the surgical area of the mouse head with 70% ethanol and iodine, a small incision was made on the skin to allow for alignment of the skull and drilling of the injection holes for 6-OHDA and virus injections. Chronic striatal DA denervation was produced through unilateral injection of 6-OHDA in the medial forebrain bundle (MFB). The toxin 6-OHDA hydrochloride (Sigma-Aldrich, Portugal) was dissolved in 0.02% ice-cold L-ascorbic acid/saline (3.2 µg free-base 6-OHDA/µL), and 1 µL was injected into the MFB (coordinates: AP= - 0.7, ML= - 1.2, DV= - 4.7), using a capillary attached to a Nanojet II Injector (Drummond Scientific, USA) at a rate of 4.6 nL per pulse every 5 s. The capillary was left in place for 2 minutes before and 10 minutes after the injection. After injecting the toxin, the capillary was replaced with a new one to inject the virus, which was injected into the striatum ipsilateral to the lesion. Each animal was then unilaterally injected with 600 nL of AAV5.CAG.Flex.GCaMP6f.WPRE.SV40 (University of Pennsylvania Vector Core) into the right dorsal striatum (AP= + 0.5, ML= - 2.3, DV= 2.3). After the surgery, the wound was closed with tissue glue (Vetbond tissue adhesive, 3M, USA) and the animal received a subcutaneous injection of the analgesic Carprofen (5 mg/kg s.c; 10 µL/10 g body weight). To prevent dehydration, mice received a subcutaneous injection of sterile glucose-ringer acetate (0.6 mL) immediately after the surgery. During the first 2 to 3 weeks post-surgery, mice received daily subcutaneous injections of sterile glucose-ringer acetate solution (0.1 mL/10 g body weight) and dietary supplementations, as necessary. In addition, mice were kept in a warming cabinet at 27 °C to keep a constant body temperature avoiding hypothermia ^9^. Lesions were also verified at the end of the study using tyrosine hydroxylase (TH) immunohistochemistry ^9,20^, and only mice with more than 90% TH depletion were included in the study (Figure S1A).

#### Chronic lens implantation and inertial sensor holder placement

Following the same surgical procedures, 3 weeks after 6-OHDA and viral injections, a gradient index (GRIN) lens (diameter: 1 mm, length: 4 mm; Inscopix) was implanted in the right dorsolateral striatum directly above the viral injection site (AP= + 0.5, ML= - 2.3, DV= 2.3), after carefully aspirating 1.8-2 mm of the overlying cortical tissue with a 30-gauge blunt needle. Care was taken to minimize bleeding before inserting the lens. Once in place, the lens was secured to the skull using superglue and a self-curing adhesive resin cement (Super-Bond C&B Kit, SUN MEDICAL CO., LTD., Japan). A small screw was placed on the surface of the skull and a layer of resin surrounded the lens and covered the screw to increase the lens bond to the skull and to minimize motion artifacts during imaging. To the self-curing adhesive resin cement, we added the mixture of black Ortho-Jet powder and liquid acrylic resin (Lang Dental, USA) on top, and finally a tape was placed to protect the lens surface. One week after the GRIN lens implantation, the microendoscope baseplate (nVistaHD, Inscopix) was attached to the microendoscope, and placed at the best focal plane to observe neuronal structures and blood vessels when present. Once the best focus was found, the baseplate was secured with a first layer of self-curing adhesive resin cement and a second layer of black cement to permanently secure the baseplate to the head cap prior to removing the microscope and attaching a baseplate cover (Inscopix) to the baseplate. Once the baseplate was placed, a small connector or holder for the wireless IMU was cemented on the back of the baseplate. The imaging field of view was inspected and allowed to clear for several days prior to imaging and behavioral experiments. Mice were excluded prior to the collection of experimental data based on imaging quality due to bad focal plane, movement artifacts or a lack of cells.

#### Drug treatment

L-DOPA methyl ester (L-DOPA also called LD; 6 mg/kg; from Sigma Aldrich, Portugal) and the peripheral DOPA decarboxylase inhibitor benserazide-HCl (12 mg/kg; from Sigma Aldrich, Portugal) were dissolved in physiological saline (9 g/L NaCl). L-DOPA was injected intraperitoneally with a volume of 10 mL/kg body weight. Vehicle (VEH) solution corresponding to physiological saline (9 g/L NaCl) was used as control. VEH was given for 2 consecutive days during which mice were recorded for imaging and behavior, and the following day, L-DOPA was administered for 3-4 consecutive days while recorded as well for imaging and behavior.

#### Open-field experiments

Experiments were conducted in a white open-field arena (40 cm X 40 cm) with a transparent acrylic on the bottom and on the side, placed inside a sound-attenuating chamber. A video camera (Flea3, Point Grey Research) was placed on the bottom of the arena to record mouse behavior from the bottom at 30-40 frames per second (fps). Before starting the experiments, mice were habituated for 2-3 days (1 hour per mouse per day) to the head-mounted equipment by using a replica of the microendoscope with the same weight (∼ 2 g) and the actual wireless inertial sensor (∼ 1.8 g). On the day of the experiment, mice were lightly anesthetized with isoflurane to facilitate mounting (and removal) of the microendoscope and the inertial sensor. Fifteen min after initial recovery from anesthesia, mice were video-recorded and striatal activity and acceleration were simultaneously recorded for 10 min, this period being referred to as baseline (BL). After BL, mice were taken out of the arena, injected with the L-DOPA or VEH and placed in a cage close to the arena. After a period of 20 min, at the peak of the L-DOPA ^20^, the animals were placed back inside the open field arena and recorded/imaged for another 10 min (see *Timeline* Figure 1A). Note that the microendoscope and the inertial sensor were left in place along the whole session, from the BL till the end of the L-DOPA/VEH recording. Acceleration was recorded using head-mounted wireless inertial sensors with a sampling rate of 200 Hz (for inertial sensor’s details see next section). The inertial sensor was secured to a connector or holder placed on the back of the microendoscope, as described above, with a consistent alignment of its axes (see Figure 1A). We note that a subset of mice was recorded with a different alignment and axes were corrected in a pre-processing step before subsequent data analyses.

#### Wireless inertial measurement unit (IMU)

The wireless, custom-made inertial measurement units (IMU) (Champalimaud Hardware Platform; WEAR motion sensor system - https://www.cf-hw.org/harp/wear) delivers a self-centered 9-axis inertial sensor containing 3-axes for accelerometer, gyroscope and magnetometer (the latest was not used in this study). The wireless IMUs are small and light (∼1.8 g) and can sample data up to 200 Hz, with a battery allowing to record up to 4 h of data. The IMUs are connected to the computer through a base station (*HARP design*, designed by the Champalimaud Hardware Platform), which provides hardware-based synchronization. We used Bonsai *visual reactive programming* (Bonsai v2.4, https://bonsai-rx.org/), which is compatible with the WEAR motion devices, to integrate and synchronize all the data coming from different sources such as the IMUs, the camera recordings and the calcium imaging recordings. Time stamps from the IMU, the video cameras and the microendoscope were then aligned using custom MATLAB scripts.

#### One-photon calcium imaging

One-photon imaging of intracellular calcium activity was acquired at 20 fps using an nVista microendoscope [lens: 1mm diameter, 4mm length, 0.5 numerical aperture; excitation: blue light-emitting diode (LED); excitation filter: 475/10 nm, 0.24-0.6 mW/mm2; emission filter: 535/50 nm; Inscopix, Palo Alto, CA] and with LED power set at 20%–40%, and gain level 4. Resulting calcium movies, video and IMU data were analyzed as described below.

#### Measurement and quantification of body acceleration with the IMU

The percentage of time moving per session was calculated using the body acceleration (BA) component of the IMU, which accurately tracks animal movement and correlates with pixel change in video measurements ^15,21^. As shown by Alves da Silva and colleagues, the average distribution of the logarithm of total BA was bimodal, with a very low acceleration distribution corresponding to immobility periods and a high acceleration distribution corresponding to periods of mobility. As done previously in the mentioned studies, to separate between rest and moving periods, we put a threshold at the value that separates the two peaks in the BA distribution. Using this threshold, which in our case corresponded to 0.05 g (ln(0.05) = -3 a.u., Figure S2C), we calculated the percentage of time that the mice moved per session and the acceleration in the periods where the mice were strictly moving (Figure 1B).

#### Abnormal Involuntary Movements (AIMs)

Three topographic subtypes of abnormal involuntary movements (AIMs; axial, limb and orofacial) were rated for 1 min every 5 min for a total of 10 min following L-DOPA injection ^9,19,20^. Axial AIMs are twisting movements or dystonic postures of neck and upper body towards the side contralateral to the lesion; limb AIMs are circular or fluttering movements of the contralateral forelimb; orofacial AIMs include twitching of facial muscles, jaw movements, and contralateral tongue protrusion. During the test, every mouse was scored on a well-characterized severity scale that is based on the duration and persistence of each dyskinetic behavior (0 = no dyskinesia; 1 = occasional signs of dyskinesia; 2 = frequent signs of dyskinesia, present for more than 50% of the observation time; 3 = dyskinesia present during the entire observation time, but interruptible by mild external stimuli; 4 = continuous dyskinesia, not interrupted by mild external stimuli). L-DOPA, at a dose of 6 mg/kg, was administered for 3-4 days and AIM scores were rated each day/session at 20, 25 and 30 min post injection.

#### Manual annotations of dyskinesia

Manual annotations of dyskinetic movements induced by L-DOPA were made frame by frame using video data recorded with a bottom camera. The annotations were performed with the software *Python video annotator* (https://github.com/video-annotator/pythonvideoannotator), for a total of 10 min per mouse, a total of 13 mice and 1 or 2 days per mouse (depicted in Figure 2A and Movie S2). Only two types of AIMs were annotated: axial and limb. The orofacial dyskinesia, although scored *live* (as described above), was not possible to annotate due to not enough resolution (30-40Hz) and angle view of the bottom camera. The criteria to detect the beginning and end of specific axial AIMs was the following. Beginning of the annotation: i) torsion of the neck/upper trunk by almost 90 degrees (bipedal position) and ii) contralateral (to the lesion) forelimb off the floor and ipsilateral forepaw extended close to the floor; end: i) 4 paws on the floor, quadrupedal position and ii) small deviation of the neck/upper trunk of less than 60 degrees. For the limb AIMs, it was as follows: the digits of the contralateral forelimb flexed and held in a fist and close to the snout was considered the beginning of the annotation; the end of the annotation corresponded to the digits of the contralateral forelimb extended and far from the snout.

Along the whole recording period, we obtained frames where axial or limb were present and frames where axial and limb appeared at the same time. Therefore, the following three combinations of annotations were considered for the analysis: *axial alone*, *limb alone* and *axial+limb.* We observed that *limb alone* did not occur very often, when present, limb was generally combined with axial (Pearson correlation between limb and axial AIMs shows a positive correlation, data not shown).

#### Video tracking and axial bending angle extraction

We used the video recordings of the bottom camera and DeepLabCut software (https://github.com/DeepLabCut/DeepLabCut) to track 9 regions of interest: nose, tail base and tail tip, left and right hind limb, left and right forelimbs, and, in order to extract additional head posture details, the base and tip of the accelerometer antenna (depicted in Figure 1D and Movie S1). The data set for the training of the network consisted of a total of 2840 manually labeled frames selected from 81 sessions (n =12 Int, n =13 Les mice; including VEH and LD together with their corresponding BL sessions). We used a training and test fraction of 0.95 and 0.05, respectively. The trained network was used to extract the positions of all regions of interest during the entire video session resulting in a set of time series for each region (i.e., x- and y-positions and the likelihood, which is an indicator of the “quality” of the extracted position). In a post-processing step, for each region of interest, we linearly interpolated the x- and y-time series if the likelihood was less than 0.6.

Once we obtained the correct tracking of all regions of interest in all mice/all sessions, we proceeded to the extraction of the axial bending angle (θ axial) as described in ^22^. In short, the θ axial was defined as the angle generated from the intersection between 2 vectors emerging from the mid-point between the hindlimbs (*mid_HL*) to the nose and to the tail base (Figure 1D).

Based on the X and Y coordinates for the nose *(Nose(xNose, yNose)*, hindlimbs *L_HL(xL_HL, yL_HL), R_HL(xR_HL, yR_HL)* and base *Tail(xTail, yTail))*, and the *mid_HL*, we calculated the *v_*Nose*_* and *v* _*Tail*_ vectors as follow:

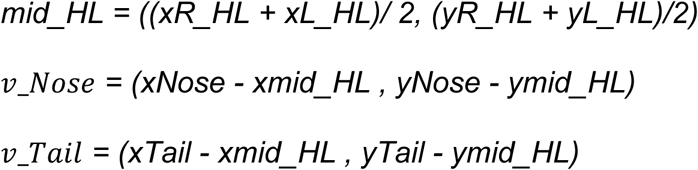

Then, to get the axial angle, θ axial, we calculated the cosine of the angle between the *“nose” and “tail”* vectors:

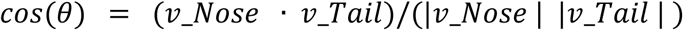

where **v**_**Nose*·*v**_**Tail**is the dot product of the two vectors and |**v**_**Nose** |and |**v**_**Tail**| are their Euclidean lengths.

Using the trained DLC network, the axial bending angle was then calculated for this cohort of mice (n = 12 intact, n = 13 lesioned) for all n = 142 sessions and the entire 10 min recordings (in total 16.9 hours).

For another group of mice that were used for calcium imaging (D1-Cre: lesioned n = 3 and intact n = 4; A2a-Cre: lesioned n = 4 and intact n = 3), we only had video-recordings from a camera placed on top of the open field that did not allow us to use the tracking method described above. We therefore developed a custom Python script to track semi-automatically the 4 body parts needed to get θ axial, i.e. the nose, tail base, left HL and right HL. We annotated these body parts every 5 to 20 frames for a total of 10 min recordings per mouse and session (n = 7 lesioned mice with 2 days per mouse, including VEH and LD; n = 7 intact mice with 2 days per mouse, LD). The calculation of the θ axial in this group of mice was done as described above.

#### Unsupervised behavioral clustering

Behavior was quantified based on the time series of acceleration and gyroscope data obtained from the IMU and the bending angle obtained with video. In short, we extracted features that capture head and body posture and motion in three dimensions. In particular, we used the following features given by the sensors: (i) total body acceleration (BA) to distinguish movement *versus* rest, (ii) gravitational acceleration along the anterior-posterior axis (GA_AP_) to extract postural changes such as those observed during rearing ^15^; (iii) gyroscope (or angular velocity) along the dorso-ventral axis that infers the head rotational behavior (θ head) and (iv) axial bending angle (θ axial), to get a measure of the axial component of dyskinesia. Example traces of each feature are shown in Figure 1C and S2A.

Total body acceleration was defined as the square root of the sum of the squares of the BA of each axis (BA= √(BA^2^_AP_ + BA^2^_ML_ + BA^2^_DV_)), where BA_AP/ML/DV_ denote the body acceleration in the anteroposterior, mediolateral, and dorsoventral axis, respectively, with respect to the animal’s head. The individual BA components were calculated by median-filtering the raw acceleration and gyroscope time series with a 7th-order one-dimensional median filter and by a subsequent Gaussian filter (type *fspecial*), to remove noise peaks. Gravitational acceleration, GA, was obtained for each axis by median and subsequent low-pass filtering (0.5 Hz cutoff, first-order Butterworth filter and a *filtfilt* filter). We used GA_AP_ to quantify postural changes (i.e., vertical head position) in the open field during resting, locomotion and rearing (as in ^15^.

Head angular velocity of the animal was obtained from the gyroscope component of the IMU data (processed as mentioned above). To capture rotational behavior, we used the dorsoventral axis and measured the cumulative head angle (θ head) from the beginning until the end of a head rotation for all rotations with a minimum of ±10 deg/sec. The value for each rotation was set to the estimated total rotation angle (Figure S2A).

The signal distribution of the above-mentioned features, BA, GA_AP_, θ head and θ axial of all mice and sessions, was used to define the thresholds for binning the feature’s time series (histogram of distributions and thresholds shown in Figure S2C). The thresholds of all features were defined as follows. The threshold for BA was set to separate moving and resting as described in *Measurement and quantification of body acceleration with the IMU.* For GA_AP_, two thresholds were defined (GA_AP_ thresholds (au) = - 0.4 and 0) to capture vertical head movements including head down, head forward and head up (including rearing). For the θ axial feature, one single threshold of 90 deg was imposed, which corresponded to the 90 deg angle of the upper torso of the mice. Finally, for the θ head feature we used a threshold of ± 25 deg to detect smaller rotations or head deviations and a higher threshold of ±165 deg to capture stronger (i.e., pathological) rotations (see Figure S3). Here, positive and negative values indicate right/ipsilateral and left/contralateral rotations, respectively. Note that the strong pathological ipsilateral rotations never happened in this study.

With the time series of the features binned using the thresholds described above, we did sliding windows of 250 ms with 80% overlap and compared them. For this, we used a similarity measure based on the so-called earth mover’s distance (EMD) ^15,32^. EMD calculates the pairwise similarity between consecutive sliding windows to get the change-points. Change-points are defined as significant changes in acceleration data corresponding to peaks detected above a threshold that was set to the average change-point value (0.005). The signal between two change-points, termed behavioral blocks or behavioral segments, had an average duration of 1.04 s (minimum duration: 100 ms, median duration: 550 ms). The obtained similarity measures were clustered using affinity propagation ^15,32^ on the EMD similarity of the behavioral blocks. Combined, for all the 600 s long recordings, this resulted in a 45,571 **×** 45,571 similarity matrix. After matching with the entire data set (see next section *Library of mouse behavior and matching procedure*), the clustering revealed a total of 56 clusters, whose signatures are shown in the *Cluster exemplars* matrix in Figure S2D. To measure the accuracy of the clustering, we did a ROC analysis, which determines whether two behavioral segments belong to the same or to different behavioral clusters based on a single threshold (Figure 1F). “True positive” corresponds to behavioral segments that were correctly classified as belonging to the same cluster whereas “false positive” corresponds to the behavioral segments from the same cluster that were misclassified as belonging to different clusters (n = 25 mice). Based on the probability of BA being below the “moving” threshold (see Figure S2A, C and D), we defined 49 out of the 56 clusters as “moving” clusters (i.e., probability of resting < 0.5) and the remaining 7 clusters as “resting” (i.e., probability of resting > 0.5).

#### Library of mouse behavior and matching procedure

Due to the computational complexity of the clustering algorithm, we created a library of mouse behavior by running the clustering for a subset of 81 sessions (n = 25 mice in total; n = 12 Int, n = 13 Les; ∼13 h of data) out of a total of 142 sessions. These included the four conditions, i.e., Int and Les, each treated with VEH and LD plus the BL sessions and one to three days per animal and treatment. We defined clusters present in a condition if they occurred a minimum of 3 s per session (corresponding to 0.5% of the session duration). Using this definition, we found 26 clusters present in Int-VEH and 23 clusters absent in Int-VEH (Figure 1G). As mentioned above, our entire data set consisted of a total number of 142 sessions, which corresponded to 16.9 h of acceleration and video data from n = 39 mice. These additional sessions were clustered on a session-by-session basis and matched to the library a posteriori using a matching procedure based on the closest EMD. We confirmed the accuracy of this approach by comparing the EMD similarity between original (library) and matched data with that of shuffled data (Figure S2B, right panel).

#### Behavioral cluster groups

For each of the 49 moving clusters, we calculated the Pearson correlation for each mouse between the occurrence of the cluster and each of the 3 combinations of dyskinesia annotations (*axial+limb*, *axial alone* and *limb alone*) and considered a correlation to be significant if the p-value was smaller than 0.05 (Figure S3A, B). We found clusters that were positively correlated with specifically one type of dyskinesia annotation, such as cluster #20, which correlated to *axial alone* in 10 out of 13 mice (Figure S3A), and others that were positively correlated with more than one type of annotation, such as cluster #7, which was correlated to *axial+limb* and *axial alone* annotations. We also found clusters that were negatively correlated to annotations (data not shown). Next, we grouped behavioral clusters at the cohort level into the following dyskinesia cluster groups: *axial+limb*, *axial alone* and *limb alone.* Due to the variability of some of the dyskinetic behavior across mice, we used the following criteria: a cluster must be significantly positively correlated to a combination of dyskinesia annotation in at least two mice. If significant negative correlations were present in some mice, the number of mice with significant positive correlations should be at least two more than the number of mice with significant negative correlations. This resulted in 9 *axial+limb* clusters, 11 *axial alone* clusters and one *limb alone* cluster (Figure 2C). In the remaining moving clusters, we found 20 clusters that were present in Int-VEH and referred to them as *other N* (*N* for “normal”). The remaining clusters appeared only in Les-LD and were largely characterized by strong pathological rotations (*path rot*). We note that behavioral clusters of the *other N* group also appeared in dyskinetic mice (Figure 2E, Les-LD). Although the behavioral signature of the clusters in *other N* Int-VEH and Les-LD was very similar, we observed differences in the probability of occurrence as shown in Figure S4B.

#### Visualization of behavioral similarity in two dimensions

For the visualization of the behavioral segments in the two-dimensional space as done previously ^15^, we used the non-linear dimensionality reduction technique t-SNE (*t-distributed stochastic neighbor embedding*, ^33^ (Figure 1H). In Figure 1H left, we show the space representation of the VEH and LD sessions of the library (‘All VEH & LD’), where each dot represents a behavioral segment and each color corresponds to a different cluster. On the right is the behavioral representation of each of the four conditions Int/VEH or LD and Les/VEH or LD. We used the matrix of pairwise EMD for all behavioral histograms (i.e., 300 ms time bins) as the input for the algorithm. We verified that the Euclidean distance in the two-dimensional projection appropriately reflected the true EMD (data not shown).

#### Classifier of *axial+limb* and *axial alone* dyskinesia

To evaluate the predictive power of the dyskinesia cluster groups, we trained a linear support vector classifier that predicts the type of dyskinesia (i.e., annotation) based on the *axial+limb* and *axial alone* cluster groups. Specifically, we used a leave-one mouse-out cross-validation approach and a fraction of 0.8 and 0.2 for training and testing, respectively. Importantly, data for training and testing was balanced by subsampling the data repeatedly and calculating the average accuracy (n = 5000). The chance level accuracy for shuffled data was calculated by random permutation of the classification labels.

#### Calcium imaging processing and analysis

Calcium recordings were pre-processed in batches using Mosaic and the *Inscopix Data Processing Software library* with a custom Matlab script for spatial binning and motion correction. Background and neuropil contamination in the one-photon microendoscopic data was corrected for using a constrained non-negative matrix factorization for microendoscopic data (CNMF-E; ^15,24,25^) and single-neuron footprints and their corresponding activities, *C_raw_*, were extracted from up to 318 neurons. In order to compare neuronal activities between BL and treatment (i.e., VEH or LD), we matched neuronal footprints within a session (i.e., when the microendoscope was kept in place) using a combination of correlation matching of the spatial footprints and manual curation.

To account for the decay dynamics of the genetically encoded calcium sensor (GCaMP6f), we quantified the event rate for each neuron by deconvolving the *C_raw_* activity traces using spike deconvolution. Since small fluctuations in the baseline fluorescence can lead to spuriously detected spikes, we used the MLspike algorithm which explicitly models a time-varying baseline ^34^. We used the following parameters for the spike detection: *τ_decay_* = 400 ms, offset correction to obtain strictly positive *C_raw_* values, multiplicative drift estimation with drift parameter 0.2, drift baseline start 0 and drift mean and standard deviation estimated for each trace by fitting a normal distribution to the distribution of offset-corrected *C_raw_*.

For the quantification of the event rate (events/sec) during the resting periods, we used the following criteria for immobility: the mouse has to have a BA lower than the threshold 0.05 g, for at least 500 ms. In the Les-LD condition, the criteria of immobility was not attained, there were only a few periods in a few mice where the mice spent from 500 to 700 ms with a BA below 0.05 g.

For the calculation of the spatiotemporal correlations, we used the same approach as in ^15^ based on the Craw time series. In short, the Pearson correlation coefficient of pairwise correlations together with the corresponding inter-neuronal distances of the spatial footprints was calculated for all neuronal pairs within an imaging session. The correlation values were binned into nine bins of logarithmically scaled spatial distances (15-750 µm) and averaged over all neurons within an imaging session. Spatiotemporal correlation plots represent the averages +/-SEM across the mice.

#### Grooming-related neurons

Since we obtained a behavioral cluster that captured grooming behavior in intact mice, we quantified the neural activity of SPNs during the *grooming* cluster group in Int-LD (Figure S6E). The average event rate was similar in D1-SPN and D2-SPN (around 0.1 Hz). We found a small percentage of *grooming-*related neurons (9-12 %), with values reached 0.35 Hz in both SPNs.

#### SPN activity during periods of non-dyskinesia

We were interested in comparing the striatal activity of intact mice under LD and Les-LD mice during periods of non-dyskinesia. We thus quantified the average event rate, the percentage of behavior-related neurons and its rate in intact mice treated with LD during the *other N* cluster group and compared it to Les-LD mice also during the *other N* (Figure S6B-D). While the average event rate in Int-LD was comparable between D1-SPN and D2-SPN (around 0.1 Hz, also comparable to the average rate in moving periods Figure 3C and E), we observed a significant increase in D1-SPN in dyskinetic mice compared to intact during *other N* behavior, while the activity was decreased in D2-SPN in dyskinetic mice (Figure S6B). The percentage of *other N*-related D1-SPNs was not significantly different between intact and dyskinetic mice, while it was significantly decreased in D2-SPNs of Les-LD mice (Figure S6C). When quantifying the neural activity of the *other N*-related neurons, we found that the D1-SPN in Int-LD had an event rate close to 0.15 Hz, which was significantly lower than the rate in Les-LD*. Other N*-related D2-SPNs however showed an activity rate of around 0.15 Hz in Int-LD, not significantly different from Les-LD mice (Figure S6D).

#### Tissue preparation and immunohistochemistry

Twenty minutes after an intraperitoneal injection of VEH or LD (6 mg/kg), mice were rapidly anesthetized with pentobarbital (500 mg/kg, i.p.; Sanofi-Aventis) and perfused transcardially with 4% (w/v) paraformaldehyde in saline phosphate buffer (PBS), pH 7.4. Brains were postfixed overnight in the same solution and stored at 4°C. Thirty micron-thick sections were cut with a Vibratome (Leica vibratome VT1000) and stored at -20°C in a solution containing 30% (v/v) ethylene glycol, 30% (v/v) glycerol, and 0.1 M sodium phosphate buffer, until they were processed for immunohistochemistry. The extent of dopamine denervation was verified in each animal by immunohistochemical staining for TH, and the transfection efficiency of the GCamP6f-GFP viral constructs was verified using an antibody that recognizes the fluorescent reporter protein GFP, according to the following protocol for double immunofluorescence. *Day 1.* Free-floating sections were rinsed in Tris-buffered saline (TBS; 0.10 M Tris and 0.14 M NaCl, pH 7.4), incubated for 5 minutes in TBS containing 3% H_2_O_2_ and 10% methanol, and then rinsed again. After 15 min incubation in TBS containing 0.2% Triton X-100, sections were blocked in serum solution, and then incubated overnight at 4°C in a mixture containing of a rabbit polyclonal α GFP primary antibody (1:1000, GFP Polyclonal Antibody, Alexa Fluor^TM^ 488, Invitrogen (Molecular Probes) Cat#A-21311), and a rabbit polyclonal α TH antibody (1:1000, Rabbit anti-TH: Peel Freez Biological, Product code: P40101-150). *Day 2.* Sections were rinsed in TBS and incubated for 45 min with the secondary antibody goat Alexa 594-coupled (1:400; Alexa Fluor 594 goat anti-rabbit, Jackson ImmunoResearch Labs Cat#115-585-045). After final rinsing steps, sections were mounted in a polyvinyl alcohol mounting medium (PVA-DABCO, Sigma Aldrich). Both placement of lens and viral transduction were confirmed using a Zeiss Axio Imager M2 fluorescence microscope.

#### Quantification of striatal dopamine depletion

Densitometry analysis of TH immunostaining on regions of interest (ROI) in the dorsal striatum were performed using the free software *NIH ImageJ 1.43.* Images were digitized using a Zeiss Axio Imager M2 microscope connected to a digital camera (Nikon DM1200F). Staining intensities were calibrated on optical density (O.D.) standards provided by the software and the average O.D. in the ROI was calculated after background subtraction. Measurements were carried out in 6 rostro-caudal sections per animal throughout the striatum. Values from the lesion side were expressed as a percentage of the mean of the values from the intact contralateral side. Animals were excluded a posteriori if the striatal TH depletion was lower than 90%.

### STATISTICAL ANALYSIS

Data analysis and statistics were performed on GraphPad Prism software, MATLAB (MathWorks) and Python. The statistical tests used and the sample size of each analysis is stated in the figure legends. The analysis of variance was performed using ANOVA (one-way or two-ways ANOVA, factorial or by repeated measures, as appropriate) followed by Bonferroni’s multiple-comparison test. For two-group comparisons, we used two-tailed Student’s t test (paired or unpaired as appropriate) or non-parametric (Mann-Whitney U or Wilcoxon signed-rank test), with Benjamini– Yekutieli procedure for multiple tests p-value correction. We used a bootstrap analysis, where indicated, for the case where we compared our data to shuffled data. Correlations were analyzed using Pearson correlation test, or the Spearman rank-order correlation test.

## Supporting information

Key ressources table

## ACKNOWLEDGEMENTS

We thank Vitor Paixão for designing and providing a modified version of the unsupervised behavioral clustering as used in Klaus et al. (2017) with the introduction of the change-point calculations. We thank Joaquim Alves da Silva for technical advice on the usage of IMUs and technical support all along this project; Catarina Carvalho and Ana Vaz for mouse colony management and mouse post-operative care; all the members of the Costa laboratory for useful discussions and support; the Histopathology Platform, Scientific Software Platform of the Champalimaud Foundation for help; the members of the Champalimaud Hardware Platform, in particular, Paulo Carriço from the hardware section for his help on developing a rotating platform for calcium imaging experiments to overcome the rotations of dyskinetic mice and Artur Silva for hardware and software assistance on the usage of the IMUs/WEAR motion sensor system; Laurent Lachaux for help with building the experimental set-ups; and Ivo Marcelo for assistance with motion correction analysis. This study was supported by the Swedish Research Council, a grant co-funded by the Marie Skłodowska Curie Actions FP7 (GS-2015-0006X) granted to C.A.; *Ayudas María Zambrano* - Alcalá University, Spanish Ministry of Universities, Next Generation UE funds to C.A.; the Swedish Research Council (2020-02696) to M.A.C.; the Swedish Brain Foundation (FO2022-0293) to M.A.C.; and the National Institute of Health funding (5U19NS104649) and the Aligning Science Across Parkinson’s (ASAP-020551) through the Michael J. Fox Foundation for Parkinson’s Research (MJFF) to R.M.C.

## AUTHOR CONTRIBUTIONS

C.A. and M.A.C. conceived the project. C.A and R.M.C. designed the experiments. C.A., A.K. and R.M.C. conceptualized the analyses. A.K., S.F.A. and C.A. configured the features and thresholds for the unsupervised behavioral clustering algorithm. C.A. and A.K. performed the behavioral clustering. C.A. performed the surgeries, one-photon calcium imaging experiments, dyskinesia scoring and annotations, and histological analysis. A.K. and C.A. performed the DeepLapCut analysis. M.M. helped organized the data and performed the analyses of average calcium imaging. A.K. and C.A. performed all other analyses. C.A., A.K., M.A.C. and R.M.C. wrote the paper. R.M.C. and M.A.C. supervised and guided all aspects of the work.

## Competing Interests

The authors declare no competing interests.

## Additional Information

Supplementary Information is available for this paper.

## Code availability

The codes will be shared in Zenodo or other repositories.

## Notes

### Competing Interest Statement

The authors have declared no competing interest.

### Summary of Updates

Author affiliations updated and distributio/resuse options updated

## REFERENCES

1. Espay, A.J., Morgante, F., Merola, A., Fasano, A., Marsili, L., Fox, S.H., Bezard, E., Picconi, B., Calabresi, P., and Lang, A.E. (2018). Levodopa-induced dyskinesia in Parkinson disease: Current and evolving concepts. Ann. Neurol. 84, 797–811. 10.1002/ana.25364.

2. Luquin, M.R., Scipioni, O., Vaamonde, J., Gershanik, O., and Obeso, J.A. (1992). Levodopa-induced dyskinesias in Parkinson’s disease: Clinical and pharmacological classification. Mov. Disord. 7, 117–124. 10.1002/mds.870070204.

3. Cenci, M.A., and Kumar, A. (2024). Cells, pathways, and models in dyskinesia research. Curr. Opin. Neurobiol. 84, 102833. 10.1016/j.conb.2023.102833.

4. Girasole, A.E., Lum, M.Y., Nathaniel, D., Bair-Marshall, C.J., Guenthner, C.J., Luo, L., Kreitzer, A.C., and Nelson, A.B. (2018). A Subpopulation of Striatal Neurons Mediates Levodopa-Induced Dyskinesia. Neuron 97, 787–795.e6. 10.1016/j.neuron.2018.01.017.

5. Parker, J.G., Marshall, J.D., Ahanonu, B., Wu, Y.-W., Kim, T.H., Grewe, B.F., Zhang, Y., Li, J.Z., Ding, J.B., Ehlers, M.D., et al. (2018). Diametric neural ensemble dynamics in parkinsonian and dyskinetic states. Nature 557, 177–182. 10.1038/s41586-018-0090-6.

6. Ryan, M.B., Bair-Marshall, C., and Nelson, A.B. (2018). Aberrant Striatal Activity in Parkinsonism and Levodopa-Induced Dyskinesia. Cell Rep. 23, 3438–3446.e5. 10.1016/j.celrep.2018.05.059.

7. Gong, S., Zheng, C., Doughty, M.L., Losos, K., Didkovsky, N., Schambra, U.B., Nowak, N.J., Joyner, A., Leblanc, G., Hatten, M.E., et al. (2003). A gene expression atlas of the central nervous system based on bacterial artificial chromosomes. Nature 425, 917–925. 10.1038/nature02033.

8. Gerfen, C.R., Engber, T.M., Mahan, L.C., Susel, Z., Chase, T.N., Monsma, F.J., and Sibley, D.R. (1990). D _1_ and D _2_ Dopamine Receptor-regulated Gene Expression of Striatonigral and Striatopallidal Neurons. Science (80-. ). 250, 1429–1432. 10.1126/science.2147780.

9. Alcacer, C., Andreoli, L., Sebastianutto, I., Jakobsson, J., Fieblinger, T., and Cenci, M.A. (2017). Chemogenetic stimulation of striatal projection neurons modulates responses to Parkinson’s disease therapy. J. Clin. Invest. 127, 720–734. 10.1172/JCI90132.

10. Albin, R.L., Young, A.B., and Penney, J.B. (1989). The functional anatomy of basal ganglia disorders. Trends Neurosci. 12, 366–375. 10.1016/0166-2236(89)90074-X.

11. Kravitz, A. V., Freeze, B.S., Parker, P.R.L., Kay, K., Thwin, M.T., Deisseroth, K., and Kreitzer, A.C. (2010). Regulation of parkinsonian motor behaviours by optogenetic control of basal ganglia circuitry. Nature 466, 622–626. 10.1038/nature09159.

12. Tecuapetla, F., Matias, S., Dugue, G.P., Mainen, Z.F., and Costa, R.M. (2014). Balanced activity in basal ganglia projection pathways is critical for contraversive movements. Nat. Commun. 5, 4315. 10.1038/ncomms5315.

13. Markowitz, J.E., Gillis, W.F., Beron, C.C., Neufeld, S.Q., Robertson, K., Bhagat, N.D., Peterson, R.E., Peterson, E., Hyun, M., Linderman, S.W., et al. (2018). The Striatum Organizes 3D Behavior via Moment-to-Moment Action Selection. Cell 174, 44–58.e17. 10.1016/j.cell.2018.04.019.

14. Cui, G., Jun, S.B., Jin, X., Pham, M.D., Vogel, S.S., Lovinger, D.M., and Costa, R.M. (2013). Concurrent activation of striatal direct and indirect pathways during action initiation. Nature 494, 238–242. 10.1038/nature11846.

15. Klaus, A., Martins, G.J., Paixao, V.B., Zhou, P., Paninski, L., and Costa, R.M. (2017). The Spatiotemporal Organization of the Striatum Encodes Action Space. Neuron 95, 1171–1180.e7. 10.1016/j.neuron.2017.08.015.

16. Sáez, M., Keifman, E., Alberquilla, S., Coll, C., Reig, R., Murer, M.G., and Moratalla, R. (2023). D2 dopamine receptors and the striatopallidal pathway modulate L-DOPA-induced dyskinesia in the mouse. Neurobiol. Dis. 186, 106278. 10.1016/j.nbd.2023.106278.

17. Cenci, M.A., and Crossman, A.R. (2018). Animal models of <SCP>l</SCP>-dopa-induced dyskinesia in Parkinson’s disease. Mov. Disord. 33, 889–899. 10.1002/mds.27337.

18. Fox, S.H., and Brotchie, J.M. (2019). Viewpoint: Developing drugs for levodopa-induced dyskinesia in <SCP>PD</SCP> : Lessons learnt, what does the future hold? Eur. J. Neurosci. 49, 399–409. 10.1111/ejn.14173.

19. Lundblad, M., Picconi, B., Lindgren, H., and Cenci, M.A. (2004). A model of l-DOPA-induced dyskinesia in 6-hydroxydopamine lesioned mice: relation to motor and cellular parameters of nigrostriatal function. Neurobiol. Dis. 16, 110–123. 10.1016/j.nbd.2004.01.007.

20. Francardo, V., Recchia, A., Popovic, N., Andersson, D., Nissbrandt, H., and Cenci, M.A. (2011). Impact of the lesion procedure on the profiles of motor impairment and molecular responsiveness to L-DOPA in the 6-hydroxydopamine mouse model of Parkinson’s disease. Neurobiol. Dis. 42, 327–340. 10.1016/j.nbd.2011.01.024.

21. da Silva, J.A., Tecuapetla, F., Paixão, V., and Costa, R.M. (2018). Dopamine neuron activity before action initiation gates and invigorates future movements. Nature 554, 244–248. 10.1038/nature25457.

22. Andreoli, L., Abbaszadeh, M., Cao, X., and Cenci, M.A. (2021). Distinct patterns of dyskinetic and dystonic features following D1 or D2 receptor stimulation in a mouse model of parkinsonism. Neurobiol. Dis. 157, 105429. 10.1016/j.nbd.2021.105429.

23. Berridge, K., and Whishaw, I. (1992). Cortex, striatum and cerebellum: control of serial order in a grooming sequence. Exp. Brain Res. 90. 10.1007/BF00227239.

24. Pnevmatikakis, E.A., Soudry, D., Gao, Y., Machado, T.A., Merel, J., Pfau, D., Reardon, T., Mu, Y., Lacefield, C., Yang, W., et al. (2016). Simultaneous Denoising, Deconvolution, and Demixing of Calcium Imaging Data. Neuron 89, 285–299. 10.1016/j.neuron.2015.11.037.

25. Zhou, P., Resendez, S.L., Rodriguez-Romaguera, J., Jimenez, J.C., Neufeld, S.Q., Giovannucci, A., Friedrich, J., Pnevmatikakis, E.A., Stuber, G.D., Hen, R., et al. (2018). Efficient and accurate extraction of in vivo calcium signals from microendoscopic video data. Elife 7. 10.7554/eLife.28728.

26. Deneux, T., Kaszas, A., Szalay, G., Katona, G., Lakner, T., Grinvald, A., Rózsa, B., and Vanzetta, I. (2016). Accurate spike estimation from noisy calcium signals for ultrafast three-dimensional imaging of large neuronal populations in vivo. Nat. Commun. 7, 12190. 10.1038/ncomms12190.

27. Maltese, M., March, J.R., Bashaw, A.G., and Tritsch, N.X. (2021). Dopamine differentially modulates the size of projection neuron ensembles in the intact and dopamine-depleted striatum. Elife 10. 10.7554/eLife.68041.

28. Lundblad, M., Andersson, M., Winkler, C., Kirik, D., Wierup, N., and Cenci, M.A. (2002). Pharmacological validation of behavioural measures of akinesia and dyskinesia in a rat model of Parkinson’s disease. Eur. J. Neurosci. 15, 120–132. 10.1046/j.0953-816x.2001.01843.x.

29. Ravenscroft, P., Chalon, S., Brotchie, J.M., and Crossman, A.R. (2004). Ropinirole versus l-DOPA effects on striatal opioid peptide precursors in a rodent model of Parkinson’s disease: implications for dyskinesia. Exp. Neurol. 185, 36–46. 10.1016/j.expneurol.2003.09.001.

30. Ryan, M.B., Girasole, A.E., Flores, A.J., Twedell, E.L., McGregor, M.M., Brakaj, R., Paletzki, R.F., Hnasko, T.S., Gerfen, C.R., and Nelson, A.B. (2024). Excessive firing of dyskinesia-associated striatal direct pathway neurons is gated by dopamine and excitatory synaptic input. Cell Rep. 43, 114483. 10.1016/j.celrep.2024.114483.

31. Barbera, G., Liang, B., Zhang, L., Gerfen, C.R., Culurciello, E., Chen, R., Li, Y., and Lin, D.-T. (2016). Spatially Compact Neural Clusters in the Dorsal Striatum Encode Locomotion Relevant Information. Neuron 92, 202–213. 10.1016/j.neuron.2016.08.037.

32. Rubner, Y. (2000). The Earth Mover’s Distance as a Metric for Image Retrieval. Int. J. Comput. Vis. 40, 99–121. 10.1023/A:1026543900054.

33. Frey, B.J., and Dueck, D. (2007). Clustering by Passing Messages Between Data Points. Science (80-. ). 315, 972–976. 10.1126/science.1136800.

34. van der Maaten, L., and Hinton, G. (2008). Visualizing data using t-SNE. J. Mach. Learn. Res. 9, 2579–2605.

