## Supplementary material for "Abnormal hyperactivity of specific striatal ensembles encodes distinct dyskinetic behaviors revealed by high-resolution clustering": Key ressources table

| REAGENT or RESOURCE | SOURCE | IDENTIFIER |
| --- | --- | --- |
| Chemicals, peptides and recombinant proteins | | |
| 6-OHDA hydrochloride | Sigma-Aldrich | H4381 |
| L-ascorbic acid, 99% | Sigma-Aldrich | A92902 |
| 3,4-Dihydroxy-L-phenylalanine | Sigma-Aldrich | D9628 |
| Benserazide hydrochloride | Sigma-Aldrich | B7283 |
| Antibodies | | |
| Rabbit anti-TH | Peel Freez Biological | P40101-150 |
| Rabbit anti-GFP Alexa Fluor-488 conjugate | Invitrogen (Molecular Probes) | Cat#A-21311, RRID: AB_221477 |
| Alexa Fluor 594 goat anti-rabbit | Jackson ImmunoResearch Labs | Cat#115-585-045, RRID: AB_2338062 |
| Bacterial and Virus Strains | | |
| AAV5.CAG.Flex.GCaMP6f.WPRE.SV40 | University of Pennsylvania Vector Core | Cat#100835-AAV5, #AV-5-PV2816 |
| Experimental Models: Organisms/Strains | | |
| D1-Cre, Tg(Drd1a-cre) FK150Gsat/Mmucd | MMRRC | #029178-UCD |
| A2a-Cre, B6.FVB(Cg)-Tg(Adora2acre) KG139Gsat/Mmucd | MMRRC | #036158-UCD |
| Softwares and Algorithms | | |
| Unsupervised behavioral clustering algorithm | This paper; Klaus et al 2017; Frey and Dueck 2007 | N/A |
| CNMF-E | Klaus et al., 2017; Pnevmatikakis et al., 2016; Friedrich et al., 2017; Zhou et al., 2018 | N/A |
| Bonsai 2.4. | Lopes et al., 2015 | https://bonsai-rx.org/; RRID:scr_017218 |
| DeepLabCut | Mathis et al., 2018 | https://github.com/DeepLabCut/DeepLabCut; RRID: SCR_021391 |
| Python Video Annotator | Champalimaud Foundation | https://github.com/video-annotator/pythonvideoannotator |
| Spyder 3.2.4 | Spyder/Python | https://www.spyder-ide.org; RRID:SCR_017585 |
| GraphPad Prism 9 | GraphPad Software | https://www.graphpad.com/; RRID: SCR_002798 |
| Inscopix Data Processing Software | Inscopix Inc. | https://www.inscopix.com/software-analysis-miniscope-imaging |
| Image J | NIH | https://imagej.nih.gov/ij/index.html; RRID: SCR_003070 |
| MATLAB | MathWorks | https://www.mathworks.com/products.html; RRID: SCR_001622 |
| Adobe Illustrator CS5 | Adobe | https://www.adobe.com/products/illustrator.html; RRID: SCR_014198 |
