## Supplemental information for "Abnormal hyperactivity of specific striatal ensembles encodes distinct dyskinetic behaviors revealed by high-resolution clustering"

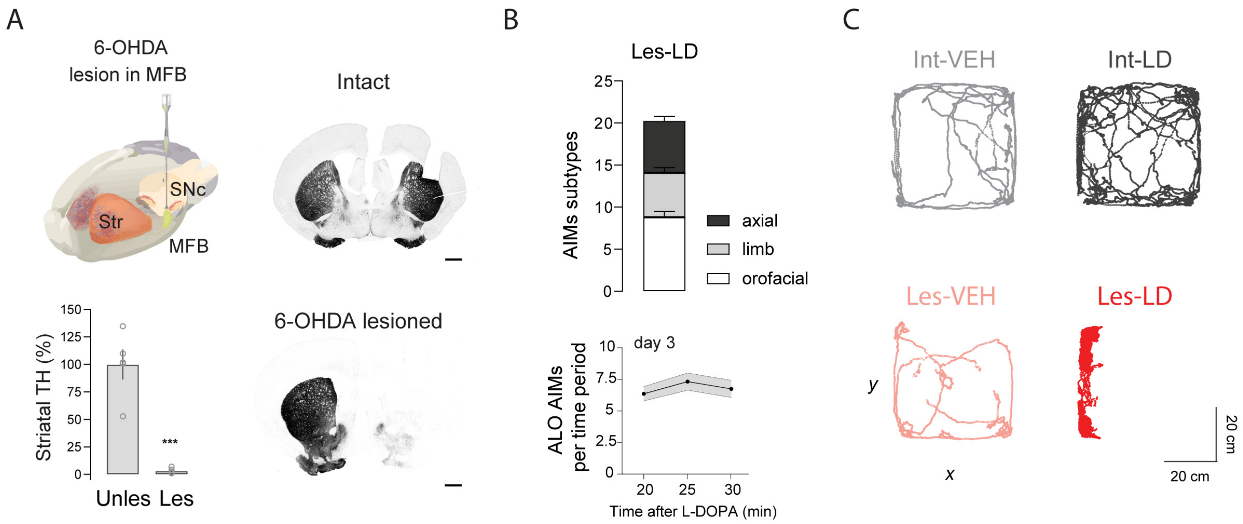


**Figure S1. 6-OHDA lesion and L-DOPA treatment induces L-DOPA-induced dyskinesia (related to Figure 1)**

(A) Top left: illustration of the unilateral 6-OHDA injection in the medial forebrain bundle (MFB) of the mouse brain. Right panels: photomicrographs of two coronal brain sections at the level of the striatum, showing a section of an intact mouse on top and a 6-OHDA lesioned mouse on the bottom (scale bar, 1mm), stained for tyrosine hydroxylase (TH), showing the lack of TH labeling in the striatum of the lesioned mouse. Bottom left: TH expression is quantified as striatal TH (%), values from the lesioned side are expressed as percentage of the mean of the unlesioned side, bars representing the mean ± SEM (n = 5 mice, STAR Methods, paired t-test, ***p < 0.001).

(B) Representation of the centroid trajectories in the open field arena of an intact mouse treated with VEH and L-DOPA (LD) (top panels), and the same for a lesioned mouse (bottom panels).

(C) Abnormal involuntary movements (AIMs) subtypes in the Les-LD condition (n = 13 mice). Bar diagrams represent the sum of AIM scores per session ± SEM, representing axial (black), limb (gray) and orofacial (white) subtypes (see STAR Methods for details of the scoring). On the right, time course of axial, limb and orofacial (ALO) AIMs per time period, on day 3 of LD treatment, rated for 1 min every 5 min for a total of 10 min following LD injection, values represent the mean ± SEM (n = 13 mice).

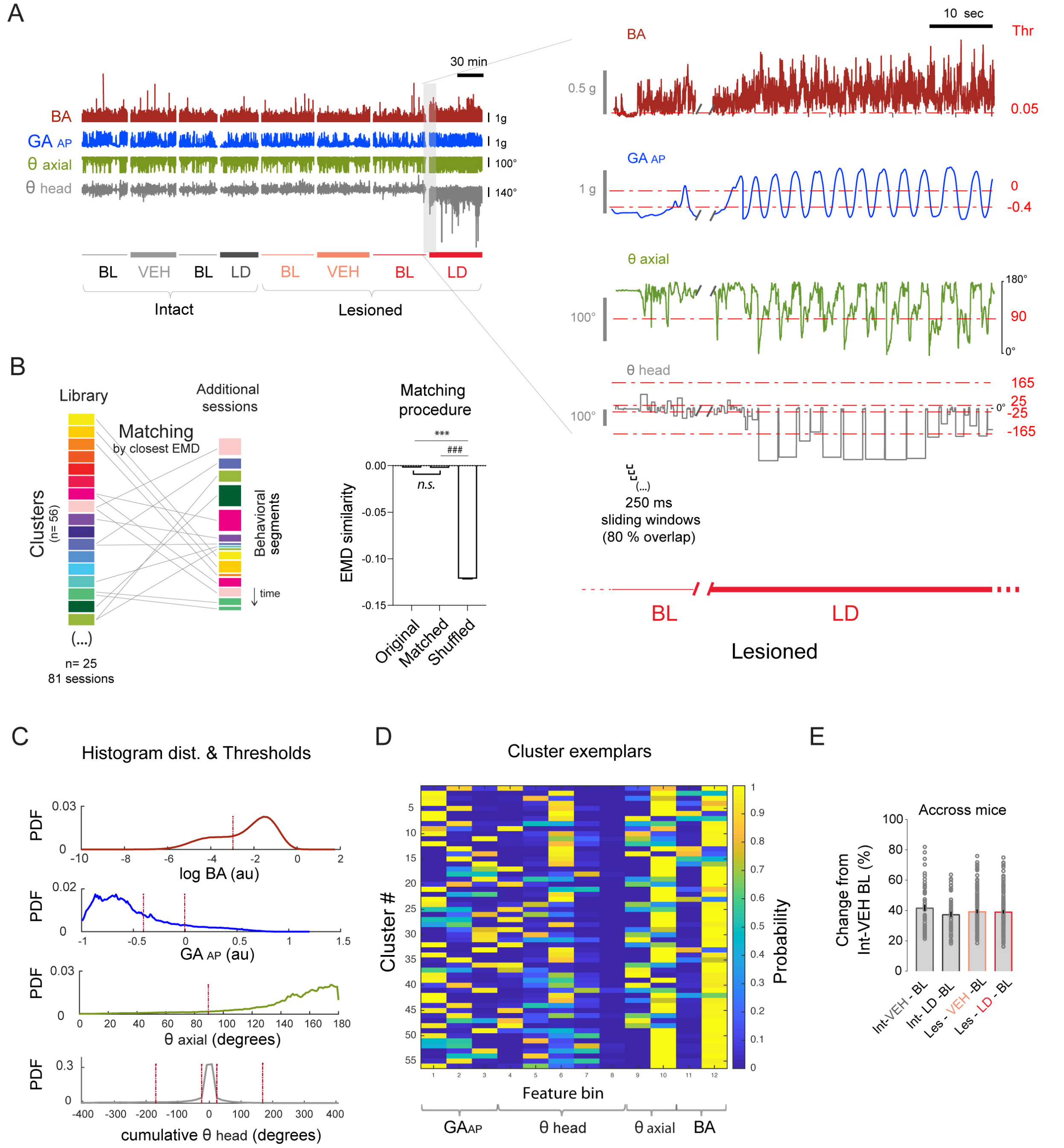


**Figure S2. Behavioral clustering algorithm details (related to Figure 1)**

(A) Example traces of the time series of the four features used for the unsupervised behavior clustering: BA (body acceleration), $GA_{AP}$ (gravitational acceleration of the antero-posterior axis), θ axial (axial bending angle) and θ head (head angle) for all four conditions, i.e., Int (n = 12) and Les (n = 13) mice, treated with VEH and LD, including the BL sessions and one to three days per animal and treatment. On the right, a zoom in the transition between Les-BL and Les-LD, showing the sliding windows of 250 ms with 80% overlap. Note the significant change in all four feature traces after LD.

(B) Additional sessions can be clustered efficiently based on matching to the reference library. Left panel shows a scheme of the matching of the behavioral segments from additional sessions to the clusters of the library (note different lengths of the behavioral segments corresponding to the different time of each behavior). The matching procedure used as a quantitative measure the Earth Mover Distance (EMD) similarity (right panel). Bar plot shows the mean of the EMD similarity ± SEM (n = 45571 behavioral segments of n = 39 mice); *Original* being the data from the library and *Matched* the data that were matched to the library (see STAR Methods). Ordinary 1-way ANOVA, $F_{(2,\ 136710)} = 55269$, $p < 0.001$. Post hoc Bonferroni's multiple comparisons test shows $^{***}p < 0.001$ original vs. shuffled; $^{\#\#\#}p < 0.001$ matched vs. shuffled; p = n.s. original vs. matched.

(C) Signal distribution of the four features, BA, $GA_{AP}$, θ axial and θ head of all mice and sessions, and thresholds for binning the feature's time series (see STAR Methods for details on the threshold's choice). For BA, a single threshold of 0.05 g (-3 in natural log scale) was used to separate moving from resting. For $GA_{AP}$, 2 thresholds were defined ($GA_{AP}$ thresholds (au) = - 0.4 and 0) to capture vertical head movements or rearing. For the θ axial feature, one single threshold was imposed (θ axial threshold = 90 deg), which corresponded to the 90 degrees angle of the upper torso of the mice (θ axial < 90 deg was considered having axial dyskinesia). Four thresholds for θ head feature (θ head thresholds (deg) = -165; -25; 25; 165). The negative thresholds corresponded to left rotations (contralateral to the lesion), -25 detects small head deviation and -165 strong pathological rotations. The positive thresholds detected right rotations (ipsilateral to the lesion), +25 detects small head deviation and +165 strong pathological rotations (note that strong ipsilateral rotations were absent in this study).

(D) Matrix showing the behavioral signature of the 56 exemplar clusters obtained. Each row represents a cluster and each column corresponds to the features' bins (obtained with the thresholds in C). As an example, BA has 1 threshold and 2 bins. That is, the

right and left bins correspond to movement and rest, respectively. Accordingly, cluster #1 is a moving cluster and cluster #6 is a resting cluster.
(E) The cluster distribution of the BL was not different in the four groups (Int-VEH/LD and Les-VEH/LD). Quantified is the change from Int-VEH baseline (BL). Comparison across mice from the BL of Int-VEH and the BL from the 4 groups (Int-VEH- BL vs. Int-VEH- BL, Int-VEH- BL vs. Int-LD- BL, Int-VEH- BL vs. Les-VEH- BL and Int-VEH- BL vs. Les-LD- BL). No significant differences were found between all four BL comparisons. Ordinary 1-way ANOVA, $F_{(3, 380)} = 1.007$, $p = 0.3895$. Post hoc Bonferroni's multiple comparisons test shows no significant difference between the four comparisons.

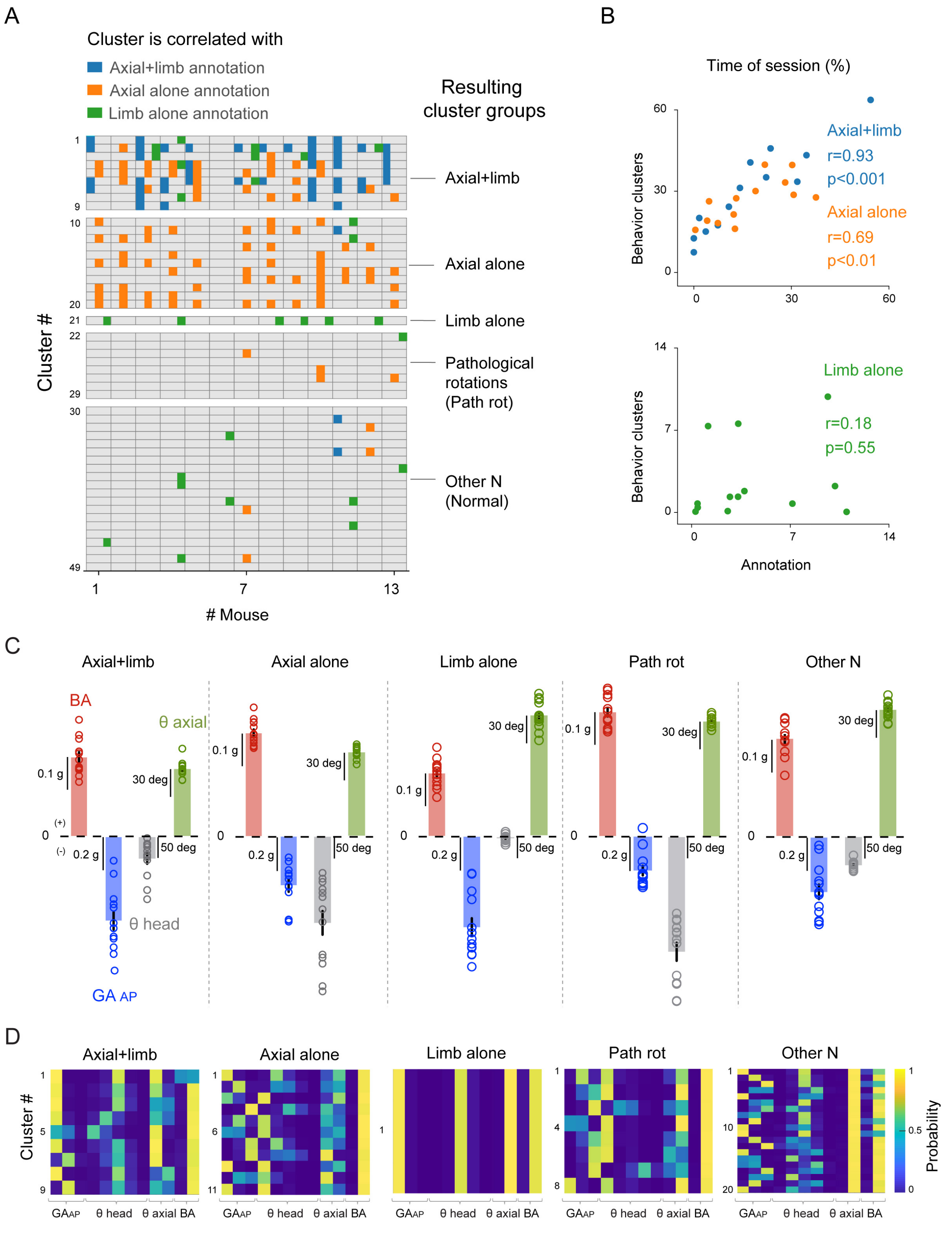

A
Cluster is correlated with
Axial+limb annotation
Axial alone annotation
Limb alone annotation
Resulting cluster groups
Axial+limb
Axial alone
Limb alone
Pathological rotations (Path rot)
Other N (Normal)
Cluster #
### Mouse
B
Time of session (%)
Behavior clusters
Axial+limb
r=0.93
p<0.001
Axial alone
r=0.69
p<0.01
Limb alone
r=0.18
p=0.55
Annotation
C
Axial+limb
Axial alone
Limb alone
Path rot
Other N
BA
θ axial
θ head
GA AP
0.1 g
0.2 g
30 deg
50 deg
D
Axial+limb
Axial alone
Limb alone
Path rot
Other N
Cluster #
GAAP
θ head
θ axial BA
Probability

**Figure S3. Cluster groups composition and features characteristics (related to Figure 2)**

(A) Map of the correlations of each behavioral cluster to the annotations, per mouse. Each row corresponds to a cluster (total number of moving clusters = 49), and each column corresponds to a mouse (n = 13 lesioned mice). The annotations are labeled in three colors, blue for *axial+limb*, orange for *axial alone* and green for *limb alone*. Per mouse, one cluster can be correlated to one annotation (example cluster #1, mouse #1, correlated to *axial+limb* annotation) or two annotations (example cluster #2, mouse #3, correlated to *axial+limb* and *limb alone* annotations). Each cluster can be correlated with one annotation (cluster #20 correlated to *axial alone* annotations), two annotations (cluster #1 correlated to *axial+limb* and *limb alone*) or three annotations (cluster #2 correlated to the 3 annotations). There are five resulting cluster groups: three groups correlated to each of the three annotations, and two groups non correlated to the annotations (*path rot* and *other N*) (see STAR Methods on the details of the cluster groups formation). Note that *limb alone* and *axial alone* are quite homogeneous cluster groups, with the majority of the clusters positively correlated to each specific annotation. In contrast, the *axial+limb* cluster group is more heterogeneous, that is, some behavioral clusters are also correlated to *axial alone* and *limb alone* annotations. Overall, this group consists of a mix of axial and limb dyskinetic components (for detailed definition of the cluster groups, see STAR Methods, 'Cluster groups').

(B) Correlation between the percentage of time spent in *axial+limb* and *axial alone* behavioral cluster groups and the percentage of time spent in the annotations. Pearson correlation test shows a significant correlation for *axial+limb*, $r = 0.93$; $p < 0.001$, and *axial alone* $r = 0.69$; $p < 0.01$; and a non-significant correlation for *limb alone*, $r = 0.18$; $p = 0.55$ (correlation plot below), indicating that the axial+limb and axial alone cluster groups truthfully capture the annotated dyskinetic behaviors.

(C) Behavioral features per cluster group. Bar plots show each feature (BA, red; GA, blue; θ head, gray and θ axial, green), values are means ± SEM (n = 13 mice). Note that the bar plots have the same scale across cluster groups to be able to visually compare magnitudes of change of the features between cluster groups.

(D) Behavioral signature of each cluster group shown per cluster. Heat maps show the behavioral signature for each of the behavioral clusters in the five cluster groups. Each row represents a cluster and each column corresponds to a bin of the behavioral feature. Note that the bins are grouped by behavior feature ($GA_{AP}$, θ head, θ axial, BA) and sorted from left to right by lower to higher numerical values, respectively. For example, the *axial+limb* behavioral clusters are associated with relatively low $GA_{AP}$ as can be seen from the high probabilities in the two left-most bins for $GA_{AP}$. In contrast, the $GA_{AP}$ component for the clusters of the *axial alone* group is more heterogeneous (i.e., clusters in this group capture the full range of $GA_{AP}$ values).

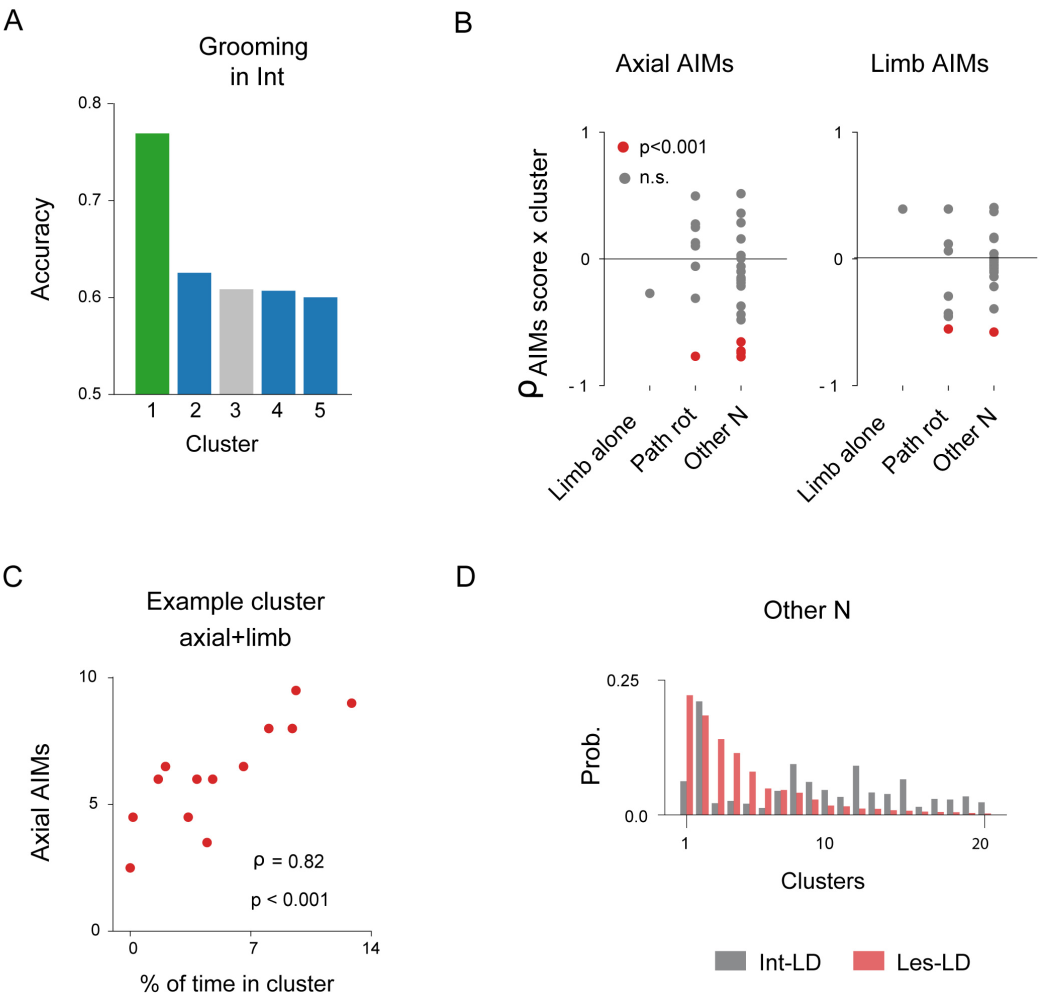


**Figure S4. Cluster groups characteristics (related to Figure 2)**

(A) Accuracy of five example clusters (*limb alone* in green, *axial+limb* in blue and *other N* in grey) to predict grooming annotations. Using a linear support vector classifier as described in STAR Methods, *limb alone* clusters predicted grooming in intact mice with an accuracy of 73 %.

(B) Correlation between the axial and limb AIMs scores and the *limb alone*, *path rot* and *other N* cluster groups. Left plot shows the correlation with the axial AIMs, the right plot with limb AIMs. Each point represents a cluster, red points being clusters significantly correlated with the AIMs scores. Note that the axial and limb AIMs are mainly negatively correlated to the *limb alone*, *path rot* and *other N* clusters, with some clusters showing significant negative correlations. Spearman rank-order correlation, red points are clusters for which the overall duration during the session was significantly correlated with the AIM scores, $p < 0.001$; gray points are non-significant correlations.

(C) Percent of time spent in some clusters is highly correlated with the axial AIMs as shown for an example *axial+limb* cluster. Spearman rank-order correlation, $p < 0.001$ (n = 13 mice, 13 sessions).

(D) Plots showing the probability of occurrence of each cluster in Int-LD and Les-LD mice in *other N* cluster group, showing its heterogeneity.

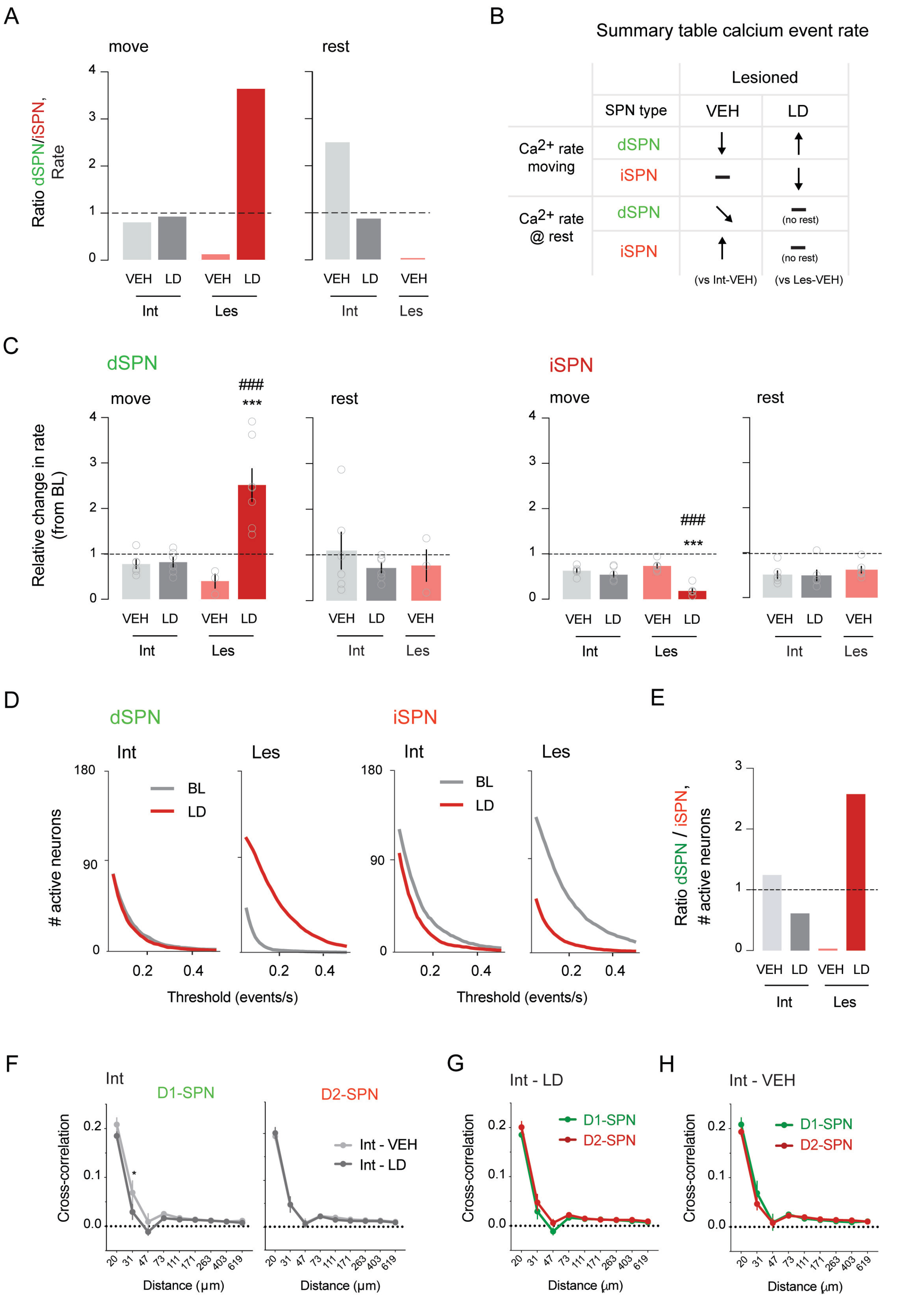
A
move
rest
Ratio dSPN/iSPN, Rate
VEH LD VEH LD
Int Les
VEH LD VEH
Int Les
B
Summary table calcium event rate
Lesioned
SPN type
VEH
LD
Ca2+ rate moving
dSPN
iSPN
Ca2+ rate @ rest
(no rest)
(vs Int-VEH)
(vs Les-VEH)
C
dSPN
iSPN
move
rest
###
***
Relative change in rate (from BL)
VEH LD VEH LD
Int Les
D
dSPN
iSPN
Int
Les
BL
LD
### active neurons
Threshold (events/s)
E
Ratio dSPN / iSPN, # active neurons
F
Int
D1-SPN
D2-SPN
Int - VEH
Int - LD
Cross-correlation
Distance (µm)
G
Int - LD
H
Int - VEH

**Figure S5. D1-SPNs and D2-SPNs modulation by L-DOPA (related to Figure 3)**

(A) Ratio of the event rate between D1-SPN and D2-SPN during moving and resting periods. Dashed line represents no changes in the ratio.

(B) Summary table of calcium event rate changes in D1-SPN and D2-SPN in lesioned mice, when moving and at rest.

(C) Relative change in event rate from BL in D1-SPN and D2-SPN during move and at rest. The plots show the relative change in the event rate from BL periods ± SEM (n = 3-7 mice per group and 6 to 18 sessions per group). For D1-SPN in move periods, ordinary 1-way ANOVA, $F_{(3,\ 18)} = 15.43$, $p < 0.0001$. Post hoc Bonferroni's multiple comparisons test shows ***$p < 0.001$ Les-LD vs. Les-VEH; $^{###}p < 0.001$ Les-LD vs. Int LD. For D1-SPN in resting periods, ordinary 1-way ANOVA, $F_{(2,\ 11)} = 0.01439$, $p = 0.9857$. Post hoc Bonferroni's multiple comparisons test shows no significant differences between groups. For D2-SPN in moving periods, ordinary 1-way ANOVA, $F_{(3,\ 20)} = 21.33$, $p < 0.0001$. Post hoc Bonferroni's multiple comparisons test shows ***$p < 0.001$ Les-LD vs. Les-VEH; $^{###}p < 0.001$ Les-LD vs. Int LD. For D2-SPN at rest, ordinary 1-way ANOVA, $F_{(2,\ 15)} = 0.5547$, $p = 0.5856$. Post hoc Bonferroni's multiple comparisons test shows no significant differences between groups.

(D) Number of D1-SPNs (left) and D2-SPN (right) at BL and after LD in Int and Les mice. The plots show the number of neurons represented as a function of the threshold (events/s) applied. Note the overlap of the curves in Int in D1-SPN and very similar profile in D2-SPN and the increase and decrease after LD in Les in D1-SPN and D2-SPN, respectively. STATS?

(E) Ratio of the number of active neurons in D1-SPN over D2-SPN during moving periods. Dashed line represents no changes in the ratio.

(F) Spatiotemporal cross-correlation between pairs of active SPNs of Int mice after VEH vs. LD. For D1-SPNs, two-way repeated measures ANOVA shows an effect of the *distance*, $F_{(8,80)} = 124.1$, $p < 0.001$; no effect of the *treatment*, $F_{(1,10)} = 2.314$, $p = 0.16$; and no effect of the interaction, $F_{(8,\ 80)} = 1.549$, $p = 0.15$. Post hoc Bonferroni's multiple comparisons test shows *$p < 0.05$ at 31 μm distance between VEH and LD. For D2-SPNs, two-way repeated measures ANOVA shows an effect of the *distance*, $F_{(8,80)} = 233.6$, $p < 0.001$; no effect of the *treatment*, $F_{(1,10)} = 0.02$ , $p = 0.89$; and no effect of the interaction, $F_{(8,\ 80)} = 0.19$, $p > 0.99$. Post hoc Bonferroni's multiple comparisons test shows no significant difference at any distance between D1-SPN and D2-SPN.

(G) Comparison of the spatiotemporal cross-correlation between pairs of active D1-SPNs vs. D2-SPN of Int mice after LD. Two-way repeated measures ANOVA shows an effect of the *distance*, $F_{(8,80)} = 198.3$, $p < 0.001$; no effect of the *treatment*, $F_{(1,10)} = 1.705$, $p = 0.22$; and no effect of the interaction, $F_{(8,80)} = 0.829$, $p = 0.58$. Post hoc Bonferroni's multiple comparisons test shows no significant difference at any distance between D1-SPN and D2-SPN.

(H) Comparison of the spatiotemporal cross-correlation between pairs of active D1-SPNs vs. D2-SPN of Int mice after LD. Two-way repeated measures ANOVA shows an effect of the *distance*, $F_{(8,80)} = 137.4$, $p < 0.001$; no effect of the *treatment*, $F_{(1,10)} = 0.179$, $p = 0.68$; and no effect of the interaction, $F_{(8,80)} = 0.727$, $p = 0.67$. Post hoc Bonferroni's multiple comparisons test shows no significant difference at any distance between D1-SPN and D2-SPN.

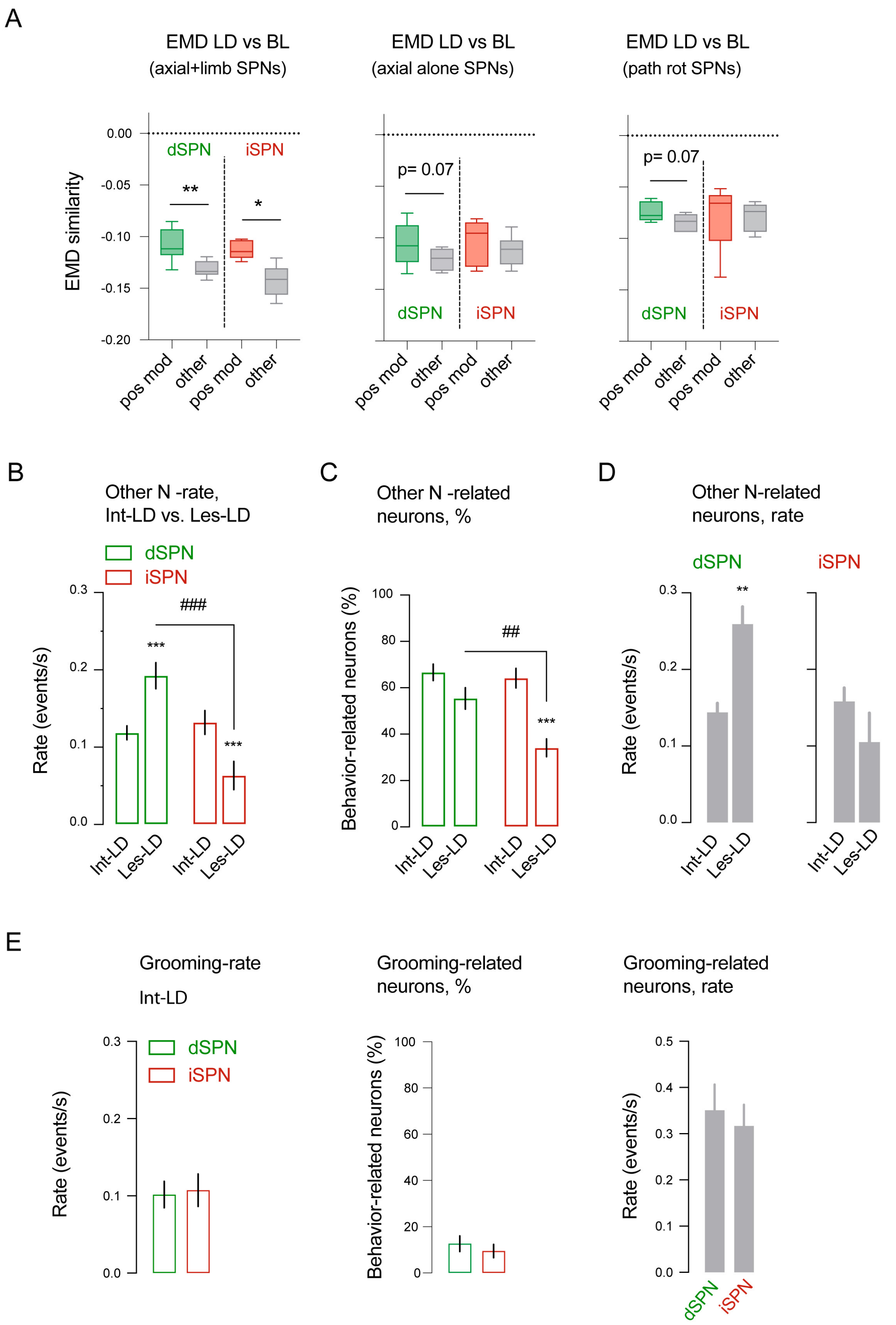
A
EMD LD vs BL
(axial+limb SPNs)
EMD LD vs BL
(axial alone SPNs)
EMD LD vs BL
(path rot SPNs)
EMD similarity
0.00
-0.05
-0.10
-0.15
-0.20
dSPN
iSPN
**
*
p= 0.07
p= 0.07
pos mod
other
B
Other N -rate,
Int-LD vs. Les-LD
dSPN
iSPN
###
***
Rate (events/s)
0.3
0.2
0.1
0.0
Int-LD
Les-LD
C
Other N -related
neurons, %
Behavior-related neurons (%)
100
80
60
40
20
0
##
D
Other N-related
neurons, rate
E
Grooming-rate
Int-LD
Grooming-related
neurons, %
Grooming-related
neurons, rate
0.5
0.4

**Figure S6. Neural activity in other N/grooming cluster groups**

(A) Comparison of the behavior EMD similarity during dyskinesia (Les-LD) versus baseline (Les-BL) for behavioral clusters that are associated with neurons in the following groups: *axial+limb*, *axial alone* and *path rot* SPNs. Specifically, for each SPN, we identified the significant dyskinesia clusters (Les-LD) and compared their EMD similarity to *other N* clusters (Les-BL) for which the neuron was positively modulated ("pos mod") vs. the remaining *other N* clusters ("other"). Box and whiskers diagrams show the values across n = 7 mice for D1-SPNs and n = 6 mice for D2-SPNs. Paired t-test shows significant differences between the EMD similarity of the positively modulated *axial+limb* D1-SPN vs. 'other' (** $p < 0.001$) and the 'pos mod' D2-SPN vs. 'other' (* $p < 0.05$). No significant differences were found in *axial alone* or *path rot*.

(B) Average rate of SPNs in Int-LD vs. Les-LD mice in *other N* cluster group. The bar plots represent the mean ± SEM (n = 6-7 mice, 2-3 sessions per mouse) of D1-SPNs (green) and D2-SPNs (red) event rates (events/s) in *other N* cluster group, comparing Int-LD vs. Les-LD. Repeated measures mixed-effect analysis: *SPN type*, $F_{(1, 11)} = 7.323$, $p < 0.05$; *lesion*, $F_{(1,10)} = 0.421$ , $p = 0.53$; interaction, $F_{(1, 10)} = 64.46$, $p < 0.001$. Post hoc Bonferroni's multiple comparisons test shows ***$p < 0.001$ Int-LD vs. Les-LD in D1-SPN; ***$p < 0.001$ Int-LD vs. Les-LD in D2-SPN, and ###$p = 0.003$ D1-SPN vs. D2-SPN in Les-LD.

(C) Behavior-related SPNs in Int-LD vs. Les-LD mice in *other N* cluster group. Bar plots represent the mean ± SEM (n = 6-7 mice, 2-3 sessions per mouse) of the percentage of neurons significantly modulated in *other N* cluster group, comparing Int-LD vs. Les-LD. Repeated measures mixed-effect analysis: *SPN type*, $F_{(1, 21)} = 8.201$, $p = 0.009$; *lesion*, $F_{(1,21)} = 24.70$, $p < 0.001$ ; interaction, $F_{(1, 21)} = 5.115$, $p = 0.03$. Post hoc Bonferroni's multiple comparisons test shows ***$p < 0.001$ Int-LD vs. Les-LD in D2-SPN; and ##$p < 0.001$ D1-SPN vs. D2-SPN in Les-LD.

(D) Event rate of *other N*-related SPNs in Int-LD vs. Les-LD in *other N* cluster group. Each bar represents the average event rate of *other N*-related neurons for *other N* cluster group, in Int-LD vs. Les-LD. Unpaired t-test shows a significant difference from Int-LD and Les-LD in D1-SPN, **$p < 0.01$; and no difference in D2-SPN.

(E) Neural activity during grooming in Int-LD mice. Left: average activity of D1-SPN and D2-SPN in Int-LD mice during *grooming* cluster group. Bar plots represent the mean ± SEM (n = 6 mice, 2 sessions per mouse). Unpaired t-test shows no significant difference between D1-SPN and D2-SPN. Middle: percentage of behavior-related neurons correlated to *grooming* cluster group. Bar plots represent the mean ± SEM (n = 6 mice, 2 sessions per mouse). Unpaired t-test shows no significant difference between D1-SPN and D2-SPN. Right: event rate of *grooming*-related SPNs in Int-LD in *grooming* cluster group. Bar plots represent the mean ± SEM (n = 6 mice, 2 sessions per mouse). Unpaired t-test shows no significant difference between D1-SPN and D2-SPN.
